# Selective direct motor cortical influence during naturalistic climbing in mice

**DOI:** 10.1101/2023.06.18.545509

**Authors:** Natalie Koh, Zhengyu Ma, Abhishek Sarup, Amy C. Kristl, Mark Agrios, Margaret Young, Andrew Miri

## Abstract

It remains poorly resolved when and how motor cortical output directly influences limb muscle activity through descending projections, which impedes mechanistic understanding of motor control. Here we addressed this in mice performing an ethologically inspired climbing behavior. We quantified the direct influence of forelimb primary motor cortex (caudal forelimb area, CFA) on muscles across the muscle activity states expressed during climbing. We found that CFA instructs muscle activity pattern by selectively activating certain muscles, while less frequently activating or suppressing their antagonists. From Neuropixels recordings, we identified linear combinations (components) of motor cortical activity that covary with these effects. These components differ partially from those that covary with muscle activity and differ almost completely from those that covary with kinematics. Collectively, our results reveal an instructive direct motor cortical influence on limb muscles that is selective within a motor behavior and reliant on a distinct neural activity subspace.

## Introduction

Motor cortex and its descending projections have expanded in certain mammalian lineages, seemingly because of the fitness conferred by the motor performance they support^1–3^. Without normal motor cortical output, certain types of movement cannot be executed^4–6^. Many other types are slower, less agile, and less effective, especially when dexterity is challenged or movements must adapt during execution^7–12^. Yet it remains poorly resolved when and how motor cortical output directly influences muscle activity through its descending projections to mediate this influence. The consequent ambiguity of direct motor cortical influence on muscles has stymied the development of more mechanistic models of descending motor control^13^.

Deficits from lesion and other inactivation of motor cortex have not clearly resolved its involvement in movement execution. Because motor cortex is involved in motor learning^14,15^ and movement preparation/initiation^16,17^, deficits could reflect disturbance to these processes on which execution depends, rather than on execution itself. Moreover, recent results indicate that motor cortical influence on muscle activity at the shortest latencies (10-20 ms in mice) differs from its influence on even slightly longer timescales (∼50 ms)^18^.

During tasks requiring motor cortex, existing results leave open several basic possibilities for the form direct motor cortical influence on muscles could take. First, motor cortex could drive the entirety of limb muscle activity patterns, with substantial compensation provided by other motor system regions following motor cortical disturbance. For example, when motor cortex needs to generate some muscle activity patterns that cannot be achieved by other regions^19^, it may assume all of the pattern-generating burden^20^. Second, motor cortex could participate in unison with the rest of the motor system in generating motor output, without playing a necessary role in determining its pattern. Here, loss of direct motor cortical influence on muscles would cause at most a non-specific, fractional attenuation of motor output. Third, motor cortex could selectively influence particular components of muscle activity, such that it informs (“instructs”) ongoing muscle activity pattern and acts distinctly from the rest of the motor system. Loss of direct motor cortical influence would then cause changes in muscle activity that themselves vary as the state of muscle activity changes.

This ambiguity about the form of direct motor cortical influence on muscles has prevented resolution of other key issues related to the mechanisms of this influence. It remains unclear whether, on balance, motor cortical output directly excites the activity of individual limb muscles, or excites and inhibits their activity depending on context. Motor cortex is thought to drive online movement corrections and the adaptation of movements based on context^9,21–24^; such a role could involve the activation and deactivation of individual muscles at different times to steer movement as context requires.

It also remains unclear what components of motor cortical output drive muscle activity. Previous descriptions of motor cortical activity have focused on components that covary with limb muscle activity^25,26^, or movement parameters like joint angles or reach direction (kinematics)^27,28^. However, if motor cortical output does not contribute to all muscle activity patterns but instead selectively alters them, we might expect that the components of motor cortical output that drive muscle activity may not reflect muscle activity in total, but only some fraction of it. Moreover, motor cortical activity that covaries with muscle activity or kinematics in total may be a consequence of monitoring or predicting body state^29^, perhaps to subserve aspects of motor control apart from directly driving muscle activation^30^. In line with this, muscle activity can be decoded from motor cortical activity during movements where this activity does not directly drive muscles^18^. Thus, the components of motor cortical activity responsible for its direct influence may differ from those to which functional significance has previously been imputed^31–35^.

### A naturalistic climbing paradigm

We sought to address these basic questions regarding direct motor cortical influence on muscles^36^. Because previous studies implicate motor cortex in adaptive limb movements in response to unpredictable sensory information^10,13,37^, we developed a behavioral paradigm that emphasizes such movements. Inspired by the natural movement repertoire of mice, we developed a paradigm in which head-fixed mice climb across a series of handholds that extend radially from a wheel, thereby rotating the wheel (Fig. 1a-d; Extended Data Fig. 1; Supplementary Movie). After each handhold accessible to the right limbs rotates 180° past the mouse, a linear actuator embedded within the wheel moves the handhold to a new, randomly chosen mediolateral position; the left handholds remain fixed (Fig. 1e; Extended Data Fig. 1b-d). This ensures that the sequence of right handholds the mouse climbs across is unpredictable (Fig. 1f), so sensory information must be used in real time to steer right limb movement. In this paradigm, water-restricted mice earn water rewards by climbing intermittently in bouts throughout hour-long daily sessions. The variation in the mediolateral position of the right handholds leads to a variation in the direction of right forelimb reaches (Supplementary Movie). A broad range of body postures are expressed (Extended Data Fig. 1e).

**Figure 1.**
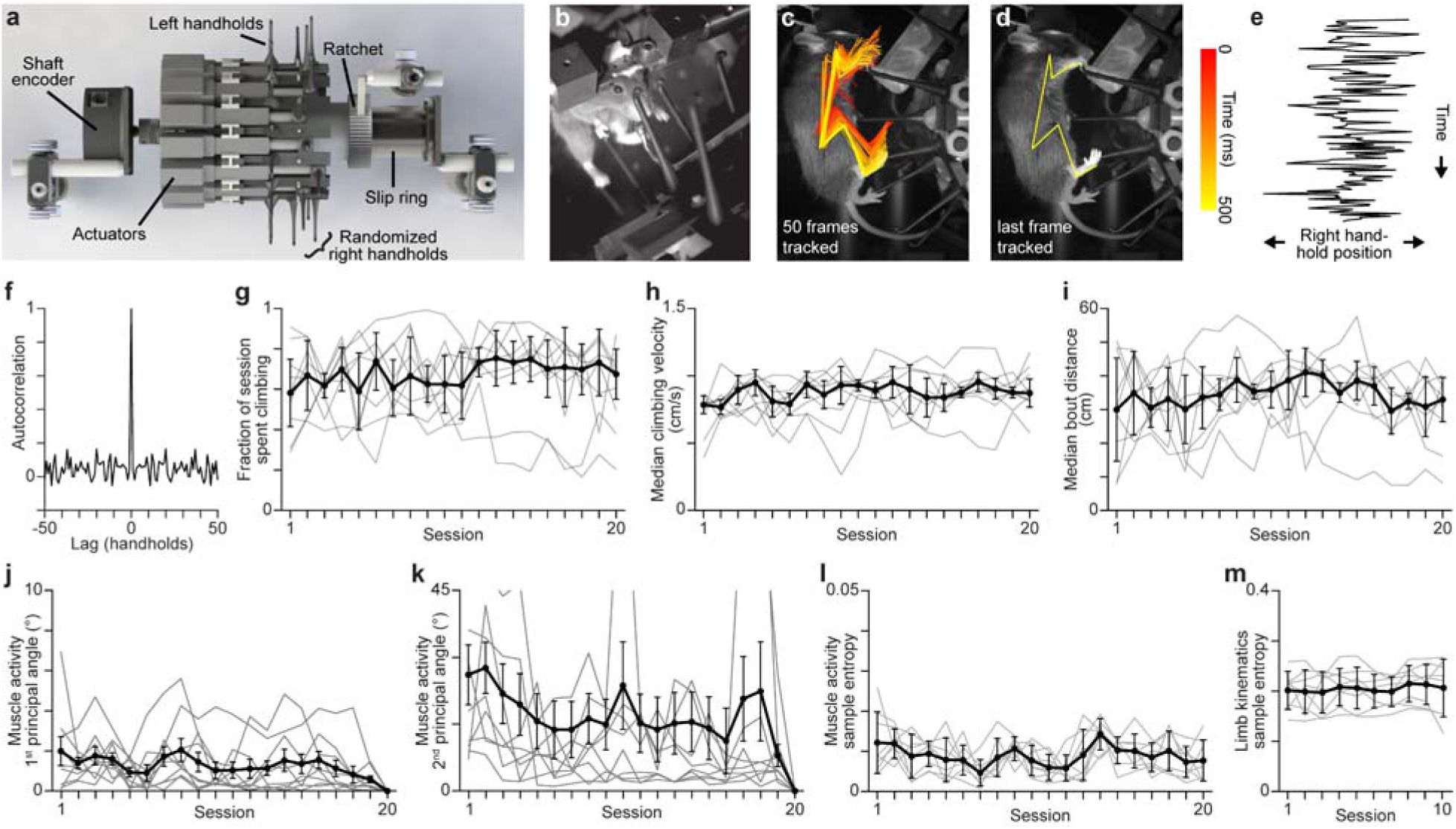
Head-fixed climbing paradigm. **a**, Bird’s-eye view of wheel apparatus for climbing. A shaft encoder measures the wheel’s angular position. Actuators randomize the position of each right handhold when they reach a point 180° away from the mouse. A ratchet ensures the wheel only rotates in one direction. A slip ring commutes voltage signals to and from the actuators. **b**, A head-fixed mouse climbing in the paradigm. **c**, Frame of side-view video of a mouse climbing, with line plots connecting points tracked on the right forelimb and hindlimb from 50 sequential images (100 Hz) overlaid. Line plot color reflects time in the sequence. Points tracked were on the shoulder, elbow, wrist, last digit of the hand, hip, knee, ankle, and edge of the foot. **d**, Same as **c**, but showing only the last frame in the sequence. **e**, Example sequence of right handhold positions over time, illustrating randomization. **f**, Autocorrelation of right handhold positions. **g**-**i**, 1^st^, 2^nd^, and 3^rd^ quartiles (n = 9 mice) for the fraction of time spent climbing (**g**), median climbing velocity (**h**), and median climbing bout distance (**i**) across sessions. Gray lines in **g**-**m** are for individual animals. Session 1 indicates the first session after mice had learned the pairing between climbing and reward, when reward dispensation switched from experimenter- to computer-controlled. **j**,**k**, 1^st^, 2^nd^, and 3^rd^ quartiles for the 1^st^ (**j**) and 2^nd^ (**k**) principal angles for EMG time series collected during each of the first 20 climbing sessions versus the 20^th^ climbing session. **l**,**m**, 1^st^, 2^nd^, and 3^rd^ quartiles for the sample entropy of muscle activity (**l**) and limb kinematics (**m**) time series during each of the first 20 climbing sessions. For each session, we took the mean across sample entropy values for each muscle, or for the X and Y positions of each tracked limb point. Sample entropy measures regularity in time series^41^.

Because it may be relevant to motor cortical involvement^38^, we assessed how the climbing mice perform varies across daily sessions. To look for progressive improvement in performance, we examined measures of bout length and climbing speed, since the reward scheme depends on them. We found that after mice are acclimated to head fixation (2 sessions) and taught the pairing between climbing and reward (1-3 sessions), there was little change on average in the time spent climbing (Fig. 1g), the velocity of climbing (Fig. 1h), and the distance of climbing bouts (Fig. 1i). To assess whether forelimb muscle activity patterns change progressively across sessions, we computed the principal angles between the first two principal components (PCs) for the activity of four muscles in the right forelimb^39^ during each session (2 PCs by T timepoints; Extended Data Fig. 1f-h). Comparing each of the first 20 daily sessions to the 20^th^ session, we found the first principal angle was generally low, averaging less than 2° (Fig. 1j,k). Though adjacent sessions appeared more similar (see lower angles for session 19), there was little indication that increasingly distant sessions were increasingly more dissimilar, which would be expected for a progressive change in muscle activity. We also found that the stereotypy in both muscle activity and limb kinematics did not show clear signs of increasing across sessions (Fig. 1l,m; Extended Data Fig. 1i). Thus, after beginning to climb for rewards, mice do not appear to progressively develop climbing skill specific to our paradigm, nor does muscle activity appear to change progressively across sessions. These results indicate that our climbing paradigm differs from those in which subjects learn new tasks and become increasingly skillful and stereotyped with repeated training^14,40^.

### Quantifying direct motor cortical influence during climbing

We next sought to quantify direct motor cortical influence on contralateral forelimb muscles across the range of muscle activity states expressed during climbing. Such influence is not seen during treadmill walking^18^, and mice can still learn new stereotyped locomotor behaviors during split-belt treadmill adaptation after bilateral motor cortical lesion^42^. However, lesion and pharmacological inactivation of motor cortex do affect the execution of novel locomotor adaptations in mice^10,42,43^. Given the different form and predictability of the movements elicited in our climbing paradigm, motor cortical influence was unclear *a priori*.

While mice (n = 8) were actively climbing, we sporadically and briefly inactivated the left caudal forelimb area (CFA, forelimb M1+S1) at random. We used transgenic mice that express Channelrhodopsin-2 in all cortical inhibitory interneurons, applying occasional 25 ms blue light pulses that covered the surface of CFA (10 mW/mm^2^; Fig. 2a). This yields a ∼50% activity reduction across cortical layers within 7 ms, which reaches 90-95% in < 20 ms^18,44^. Light pulses were always > 4 seconds apart to allow recovery of neural activity between events; on average, ∼100-200 trials were collected during each daily session (11-37 sessions per animal). Equivalent events without blue light were notated in recordings to serve as control trials. Random trial timing ensured broad coverage of the muscle activity states each mouse expressed during climbing. We found that inactivation and control trial averages diverged ∼10 ms after light onset, which reflects the shortest latency at which CFA output influences muscles^18^ (Fig. 2a-c, Extended Data Fig. 2a-d). We also found that inactivation effects were similar in form across mice (Fig. 2a; Extended Data Fig. 2b) and strikingly consistent both within and across sessions (Extended Data Fig. 2e-g). Thus CFA directly influences muscle activity during climbing, as we previously observed in mice performing a trained forelimb reaching task^18^.

**Figure 2.**
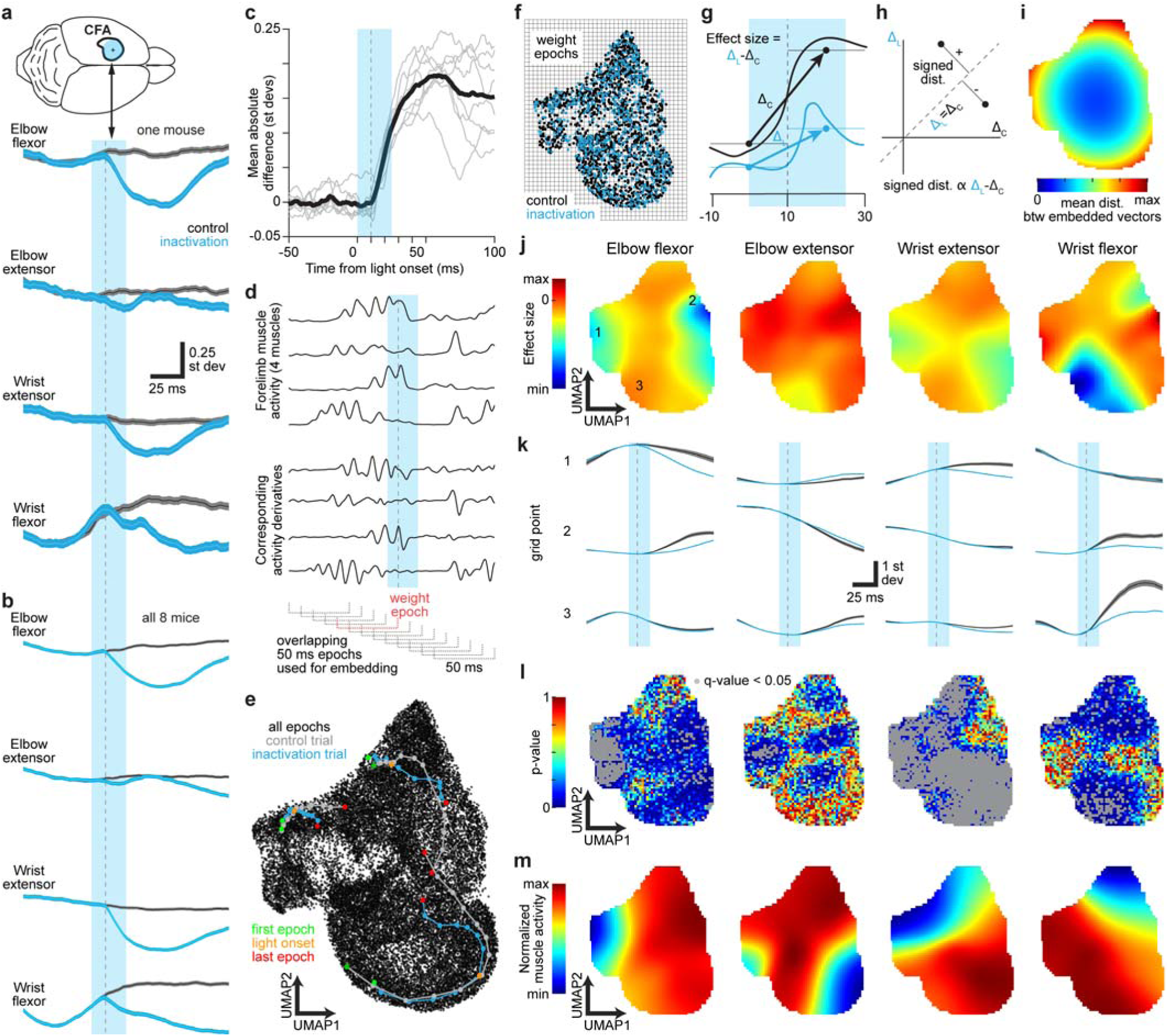
Comprehensive assessment of CFA influence across muscle activity states. **a**, Control (n = 1,671 trials) and inactivation (795 trials) trial averages (mean ± sem) for four muscles in one mouse. Vertical cyan bars in **a**-**d,g,k** indicate the 25 ms epoch of blue light applied to CFA and gray dotted lines are 10 ms after light onset. Because we have z-scored muscle activity measurements using the mean and standard deviation from each given session, here and throughout we express measurements in standard deviations of the recorded signal. **b**, Control (18,397 trials) and inactivation (9,029 trials) trial averages for all eight mice. **c**, Mean absolute difference between inactivation and control trial averages across all four muscles. Light gray lines are individual animals, and the solid black line is the mean across animals. For baseline subtraction, control trials were resampled to estimate the baseline difference expected by chance. **d**, Example of muscle activity and its corresponding first derivative surrounding trials that were used for creating muscle activity state maps. The weight epoch immediately precedes when effects begin. **e**, Example muscle activity state map from one animal. Larger, connected dots show examples of states for sequential overlapping epochs from individual trials. Pairs were chosen based on their similarity during the weight epoch. **f**, Grid overlaying a map including only points from the weight epochs used for weighting trials in grid point trial averages. **g**, Schematic of the calculation of inactivation effect at each grid point from the control (black) and inactivation (cyan) trial-averaged muscle activity. Δ_C_ and Δ_L_ reflect the slopes of lines connecting the average activity just before to just after the inactivation effect begins. **h**, Schematic illustration of the effect size on a plot of Δ_L_ versus Δ_C_. Their difference is proportional to the distance from the identity line. **i**, Map in which each grid point is colored by the mean distance, in the full 80D space, between all pairs of embedded state vectors, with each individual distance weighted by a Gaussian function of the pair’s mean distance from the grid point on the 2D map. The Gaussian function is the same as that used for inactivation maps. **j**, Inactivation effect maps for the four recorded muscles. Color scale max and min reflect the maximum and minimum effect sizes across all four muscles collectively. Panels **j**-**m** show representative results from one mouse. **k**, Grid point-averaged muscle activity from control (gray, mean ± sem) and inactivation (cyan, mean) trials, for three example grid points from the maps in **j**. **l**, Maps of p-values computed for inactivation effects at each grid point. Q-values (gray overlay) reflect the expected rate of false discovery below the corresponding p-value^46^. **m**, Maps showing the average activity for the four recorded muscles at each grid point. Color scale max and min reflect the maximum and minimum activity level for each muscle separately. Darker blue regions reflect states where the given muscle is inactive. Darker red regions reflect states where the given muscle is highly active, up to between 2.7 and 5.5 standard deviations above the mean.

To initially gauge whether direct motor cortical influence varies throughout climbing, we examined the effects of CFA inactivation during three stereotypical features of climbing – pulling a handhold down with the right forelimb, reaching the right forelimb up, and palpation of the right handhold while grasping it (Extended Data Fig. 3). We assembled trial averages for muscle activity and limb kinematic time series aligned on trial onsets that occurred during each feature. Effect magnitude appeared variable across features. Effects were also more prominent in trial-averaged muscle activity than limb kinematics across all three features, which we explore further below.

We thus more comprehensively assessed CFA influence at different muscle activity states during climbing. We first sought a means for collecting together inactivation and control trials that began at similar muscle activity states. Plotting trials according to linear functions of muscle activity at trial onset led to an uneven distribution of trials across plots (Extended Data Fig. 3d,e). This was suboptimal for efficiently utilizing the statistical power afforded by our trials to differentiate CFA influence across states (see Methods). We thus explored the use of dimensionality reduction methods that organize states according to their *n* nearest neighbors. We used UMAP^45^ to generate a 2D map of the muscle activity states expressed by each mouse where similar states are close together. To overcome EMG signal variability and account for history dependence of CFA effects, we used states spanning 50 ms epochs rather than individual time points. To ensure proximity on maps among states visited close together in time, we defined epochs that overlapped in time. Muscle activity traces surrounding each trial, together with their corresponding first derivatives, were subsampled in 5 ms increments and divided into overlapping 50 ms epochs that began every 10 ms (Fig. 2d). For each 50 ms epoch, the muscle activity and first derivative trace segments were concatenated into a single vector (8 segments x 10 time bins = 80 elements). UMAP was then applied to map vectors onto two dimensions (Fig. 2e). On the resulting maps, embedded state vectors (points) from successive epochs form trajectories that reflect the sequence of states surrounding control and inactivation trials.

To measure direct CFA influence at different muscle activity states, we quantified the immediate inactivation effects for trials beginning from states within local neighborhoods on the maps. We first defined a grid over each map (Fig. 2f). For each muscle, we computed its trial-averaged activity at each grid point, separately for inactivation and control trials. For these averages, we used all trials, but we weighted each by a Gaussian function of the Euclidean distance between the given grid point and the point for the epoch just prior to when an inactivation effect could begin on the given trial (−40 to +10 ms from trial onset, “weight epoch;” Fig. 2d). Trial weights are hence not influenced by inactivation effects. As a consequence, weight epoch states from control and inactivation trials are similarly distributed across maps (Fig. 2f). We set the Gaussian’s standard deviation to be roughly 10% of the map width, so trial averages are heavily weighted toward trials beginning at states close to the given grid point. Only grid points close to a substantial number of weight epoch states were subsequently considered (“valid grid points;” see Methods). Separately for each muscle, we measured the size of the inactivation effect at each valid grid point as the difference between the rate of change in inactivation and control trial averages from 0 to 20 ms after trial onset (Fig. 2g,h). We then plot the resulting effect sizes at each valid grid point across the map, producing an “inactivation effect map” (Fig. 2j; Extended Data Fig. 4a-d). Maps for different muscles in a given mouse show wide variation in the magnitude and sign of inactivation effects across grid points (Fig. 2j,k). We resampled from control trials to compute a p-value for each grid point’s effect size (Fig. 2l; Extended Data Fig. 4a-c). Map structure was not strongly dependent on the choice of key parameters (Extended Data Fig. 4f-i).

Because UMAP is nonlinear, it is not clear how the distance across maps will correspond to differences in muscle activity state. To address this and clarify muscle activity levels at different map locations, we plotted the average activity of each individual muscle at each grid point using the same Gaussian weighting method as above (Fig. 2m). These plots showed smooth and gradual variation across grid points. We also plotted the average similarity between nearby state vectors across maps (Fig. 2i; Extended Data Fig. 4e). The only prominent variation we observed was a gradual increase in pairwise distance from the map center to the edges; there was no indication of abrupt changes in pairwise distance. Thus muscle activity state varies smoothly across state maps at the resolution of our inactivation effect maps.

### CFA acts primarily by selectively exciting physiological flexors

We then analyzed inactivation effect maps to distinguish among the possible forms direct CFA influence on forelimb muscles could take. We first generated histograms of the p-values computed for effects on each muscle at all valid grid points. These histograms consistently showed a skew toward zero, reflective of a substantial fraction of grid points where the null hypothesis of no effect was false (Fig. 3a,b). From these distributions, we estimated the fraction of grid points showing effects (Fig. 3c). The mean estimated fractions were 0.62, 0.22, 0.73, and 0.37 for elbow flexor, elbow extensor, wrist extensor, and wrist flexor muscles, respectively. These estimates were significantly above zero for all four muscles (p = 0.008 or p = 0.016, Wilcoxon signed-rank test). Control maps generated from comparisons between separate sets of control trials yielded uniform distributions, as expected under the null hypothesis (Extended Data Fig. 5a). These results show that direct CFA influence on muscles is specific to a subset of muscle activity states.

**Figure 3.**
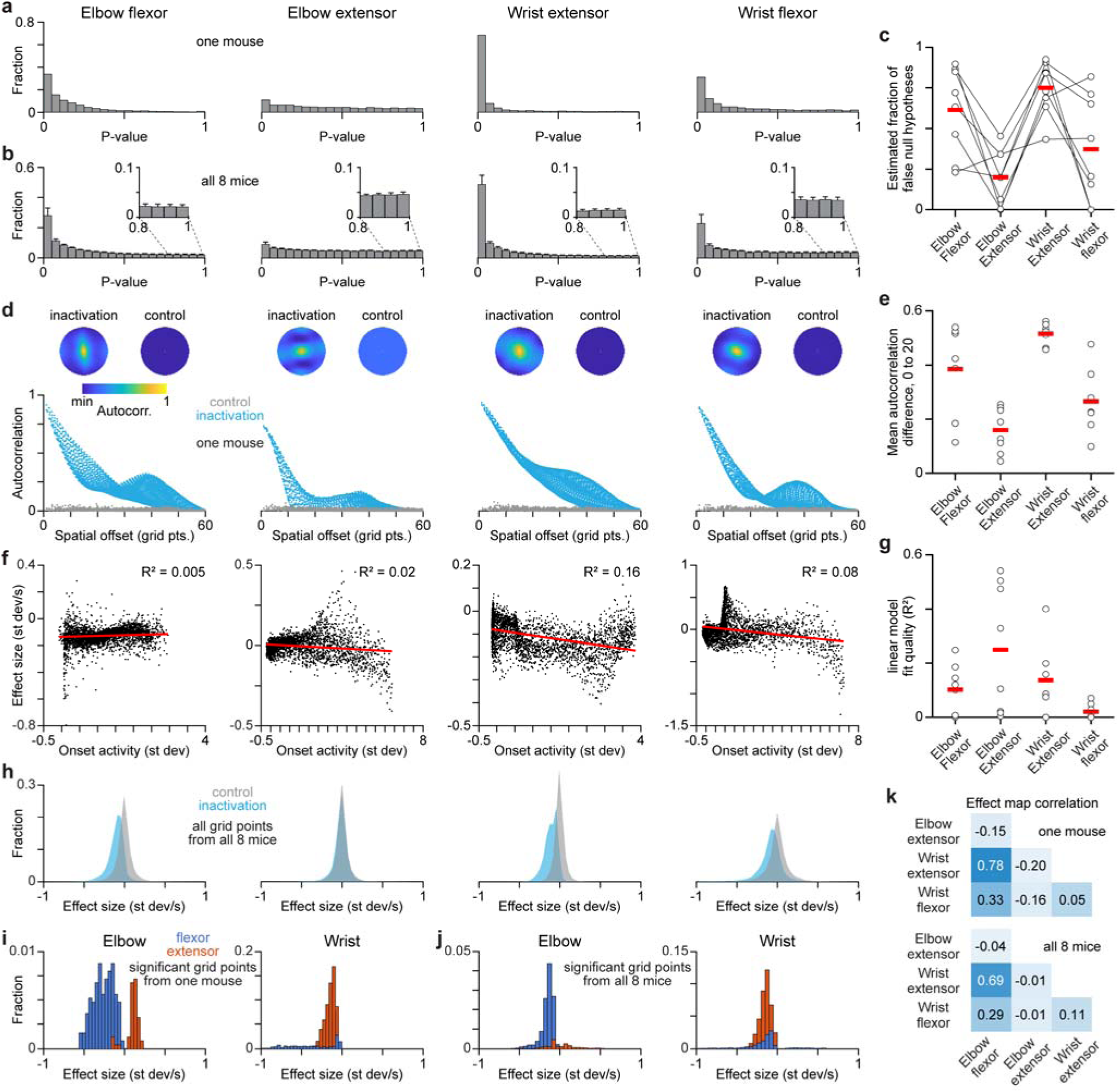
**CFA selectively excites physiological flexors. a,b**, P-value distributions for inactivation effects on each muscle for all grid points in one mouse (**a**) and across all 8 mice (**b**). **c**, Estimated fraction of grid points at which the null hypothesis of no effect is false, calculated from p-value distributions, for individual animals (black circles) and the mean across animals (red bars). **d**, For one mouse, the 2D autocorrelation for inactivation effect maps and control maps generated with only control trials (top) and scatterplots of correlation values versus their spatial offset (lag from zero offset). **e**, Difference between inactivation effect maps and control maps in their mean autocorrelation over spatial offsets from 0 to 20 grid points for individual mice (black circles), and the mean across animals (red bars). **f**, Scatterplots of inactivation effect size versus muscle activity at trial onset for one animal. Each point reflects a different grid point. R^2^ is for a linear fit (red). **g**, R^2^ for linear fits to scatterplots of inactivation effect size versus muscle activity at trial onset for individual mice (black circles), and the mean across animals (red bars). **h**, Effect size distributions for all grid points across all 8 mice, separately for inactivation effect maps and control maps. **i**,**j** Effect size distributions for all significant grid points from one mouse (**i**) and all 8 mice (**j**). **k**, 2D correlation between inactivation effect maps for different muscles for one mouse (top) and the means across all 8 mice (bottom).

We next assessed whether CFA influence varies in magnitude across muscle activity states. If this were true, then the 2D autocorrelation of inactivation effect maps should be significantly above what is expected by chance. We computed the 2D autocorrelation of both inactivation effect maps and control maps, observing substantially heightened autocorrelation in the former (Fig. 3d). To assess whether these differences were significant, we computed the mean difference between inactivation effect map autocorrelation and that of control maps, averaged over spatial lags up to 20 grid points (Fig. 3e). These differences were significantly above zero for all four muscles (p = 0.008 for each muscle, Wilcoxon signed-rank test). We also found that the magnitude of CFA influence across muscle activity states differed significantly between muscles in six out of eight mice (p < 0.004, p = 0.19 and 0.83 in two other mice, Kruskal-Wallis test). CFA influence magnitude was not simply proportional to muscle activity magnitude; R^2^ for linear fits to effect size versus muscle activity magnitude were low (Fig. 3f,g; Extended Data Fig. 5b-e) and the residuals were significantly nonuniform (p < 10^-10^ for all mice, K-S test).

A number of previous observations indirectly suggest that primary motor cortex may preferentially control certain muscle groups more so than their antagonists^47–49^. We thus compared the distributions of effect sizes across grid points for each muscle. We found larger deviations from control effect sizes for the elbow flexor and wrist extensor (Fig. 3h). The estimated fraction of grid points showing effects (false null hypotheses, Fig. 3c) was significantly greater for the elbow flexor (61% higher; p = 0.007, Wilcoxon rank-sum test) and wrist extensor (43% higher; p = 0.015), compared with their respective antagonists. This indicates that CFA output preferentially influences elbow flexors and wrist extensors, which can be grouped together as *physiological flexors* because of their coactivation during locomotion and during the flexion reflex^50^.

We also assessed whether these differences in effects on different muscles might extend to the direction of effects. Reduction or elevation of muscle activity following inactivation indicate that CFA output activates or suppresses muscle activity, respectively. We examined effect sizes that were significantly different from zero, finding that for the physiological flexors, effects were always a reduction in muscle activity (Fig. 3i,j). However, the elbow extensor exhibited both reduction and elevation, and the wrist flexor showed elevation at some states as well (Extended Data Fig. 5f). We also observed that inactivation effect maps for the physiological flexors were more highly correlated compared to those for all other pairs of muscles (Fig. 3k), suggesting a greater degree of coordinated control of these muscles. Collectively, these results indicate that CFA activates physiological flexors to varying degrees and only at some muscle activity states (i.e., influence is selective), while activating *or suppressing* their respective antagonists at a smaller fraction of states. Thus, direct CFA influence instructs muscle activity pattern during climbing.

### Weak covariation between CFA influence and gross kinematic state of the forelimb

A number of previous observations suggest that motor cortex may, to some extent, control the limb via commands that dictate its kinematics rather than muscle activation^51,52^, although this remains controversial^32,53^. If CFA were dictating contralateral forelimb kinematics, we reasoned that direct CFA influence on muscles should correlate with the orientation of the contralateral forelimb. We therefore probed for this correlation.

Here we mimicked the approach we took to assess how CFA influence covaries with muscle activity state. We computed new state maps with UMAP using vectors composed of the horizontal and vertical positions of sites on the right forelimb tracked from video (Fig. 4a,b). Nearby points on these maps thus reflect 50 ms epochs of limb kinematics that are similar. The resulting 2D maps separated states that correspond to different limb orientations into different map regions (Fig. 4c), with the cyclic changes in limb orientation during iterative climbing ordered around the map. Using these maps, we quantified the effects of CFA inactivation on each forelimb muscle at grid points covering the maps as above. Histograms of the inactivation effect sizes across all grid points showed deviations from controls (Fig. 4d). However, p-value distributions for effects on each muscle showed very limited skew toward zero, indicating discernible effects on only a small fraction of grid points (Fig. 4e). Thus, CFA influence on muscles does not covary with forelimb orientation nearly as well as it does with muscle activity state.

**Figure 4.**
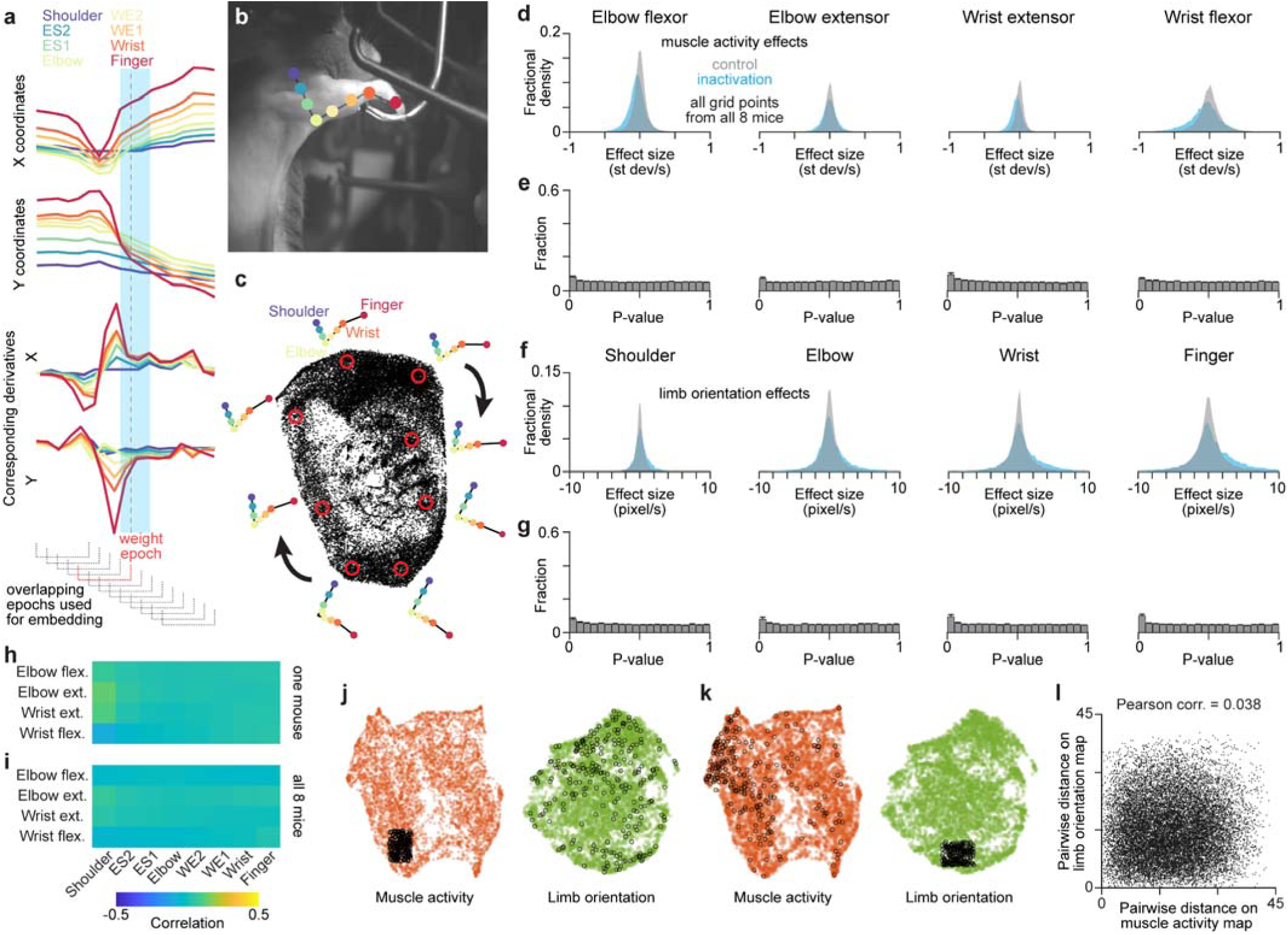
Gross forelimb kinematics capture CFA influence worse than muscle activity. **a**, Time series of tracked forelimb sites and their corresponding first derivatives surrounding inactivation and control trials are windowed into overlapping segments. Segments are used to create a 2D embedding via UMAP. **b**, Image showing the locations of the eight sites tracked on the forelimb, according to the color code in **a** (ES1,2: 1^st^ and 2^nd^ sites between elbow and shoulder joints, WE1,2: 1^st^ and 2^nd^ sites between wrist and elbow joints). **c**, Example map of forelimb orientation states from one mouse, along with the trial-averaged positions of the forelimb sites at selected grid points within the map (red circles). Because video was captured at 100 Hz, time series segments used here had 10 ms spacing between points instead of 5 ms, as in Fig. 2. **d-g**, Distributions of the sizes of inactivation effects on muscles (**d**), p-value distributions for inactivation effects on muscles (**e**), distributions of the sizes of inactivation effects on four main forelimb sites (**f**), and p-value distributions for inactivation effects on four main forelimb sites (**g**), calculated using forelimb orientation maps, across all grid points and all 8 mice. **h**,**i**, Matrices indicating the Spearman correlation between each muscle activity and limb kinematic time series for one mouse (**h**) and all eight mice (**i**). **j**,**k**, Maps of muscle activity states (left, orange) and limb orientation states (right, green) from one animal. Black circles in each panel mark corresponding sets of embedded vectors on the two maps (i.e., those coming from the same set of epochs). Both map types used the same time point spacing (10 ms) and equivalent amounts of data. **l**, For the embedded vectors from 200 randomly selected epochs, the Euclidean distance between all possible pairs of those vectors on the muscle activity map in **j**,**k** plotted against the distance between their corresponding vectors on the limb orientation map.

If CFA output dictates forelimb kinematics, we might expect that CFA inactivation would perturb limb kinematics themselves. We therefore quantified the effect of inactivation on the position of the sites tracked at the shoulder, elbow, wrist, and finger. Inactivation effects were computed as above, but using trial-averaged site position in place of trial-averaged muscle activity. Again, histograms of inactivation effect sizes did show deviation from controls (Fig. 4f), but p-value distributions showed very limited skew toward zero. This indicates there are discernible effects on only a small fraction of grid points (Fig. 4g). Collectively, these results suggest that CFA output directly specifies muscle activity and not limb orientation.

Given an expectation that muscle activity and limb kinematics should covary, these results may seem surprising. However, we note that during adaptive, nonstereotyped motor behaviors like climbing in our paradigm, linear covariation between muscle activity and limb orientation will not necessarily be strong, due to the complex causal relationship between these variables. To assess how well coupled muscle activity and limb kinematics were in our paradigm, we measured the Spearman correlation between their time series. These correlations were generally close to zero (Fig. 4h,i). Next, we made state maps using either data type from the same set of recording epochs. We found that the proximity of muscle activity states on maps only weakly predicted the proximity of the corresponding limb orientation states, and vice versa (Fig. 4j-l). Subsets of states close together on one map type corresponded to states that were widely distributed on the other map type. This decoupling between muscle activity and limb kinematics may have helped reveal that CFA influence is not well-organized by limb orientation.

### CFA firing patterns during climbing

To assess what components of CFA output might drive the direct influence we have identified, we next sought to determine how CFA firing patterns covary with CFA influence on muscles. We used Neuropixels to measure the firing of CFA neurons across cortical layers in three mice for which inactivation effect maps were computed (Fig. 5a-c). After completing collection of inactivation trials for mapping influence, we recorded neural activity in CFA with acutely-implanted Neuropixels during the next 3-4 daily behavioral sessions. The majority of both wide- and narrow-waveform units had higher firing rates during forelimb muscle activity as compared with periods when all recorded muscles were quiescent (Fig. 5d). Over 80% of recorded units had firing rate time series that were significantly correlated with the activity of at least one muscle (Fig. 5e). Fitting muscle activity time series with neuronal firing rates using ridge regression and Weiner cascade models yielded similar accuracy across muscles (Extended Data Fig. 6a).

**Figure 5.**
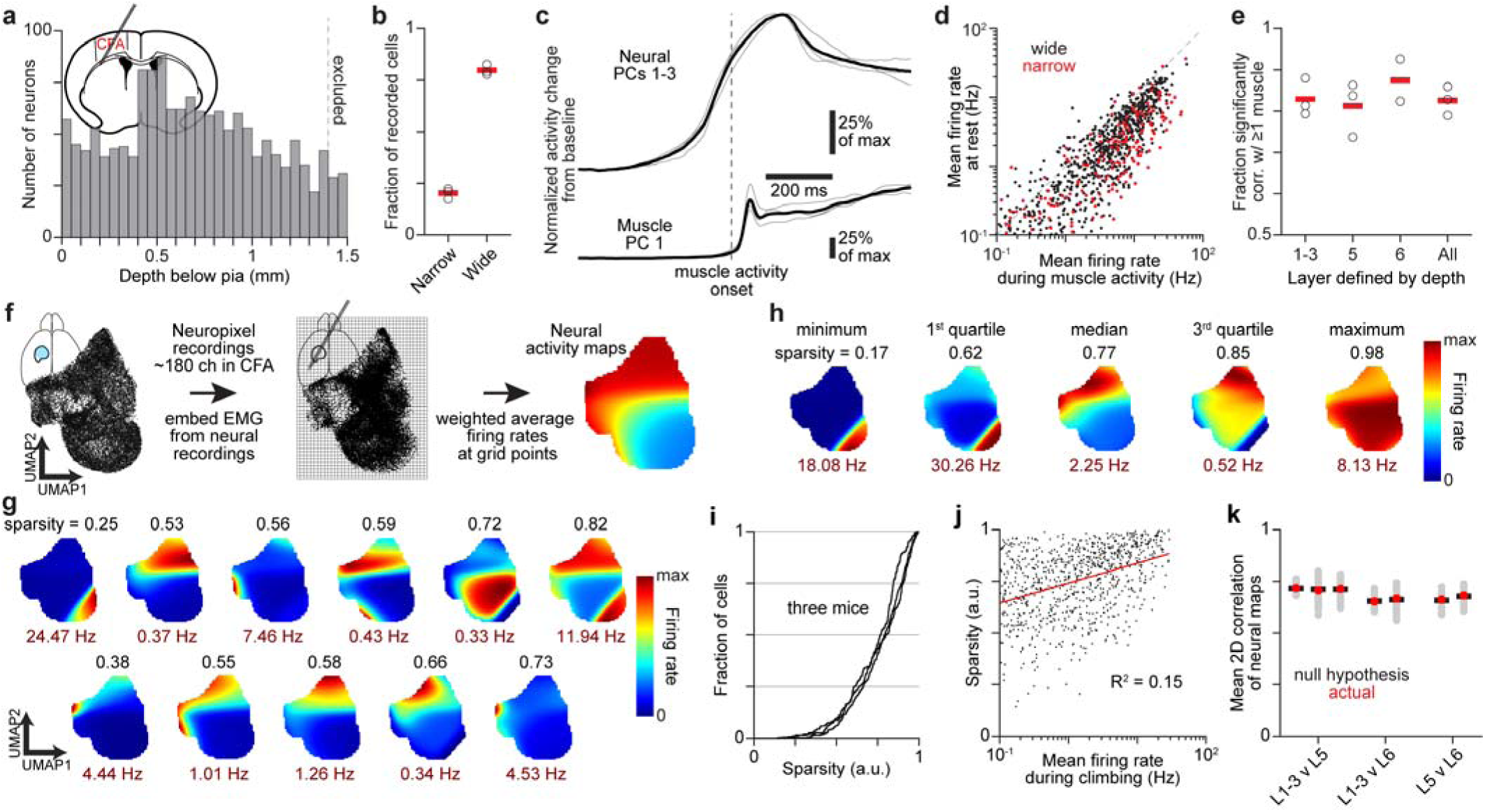
CFA firing patterns vary in their sparsity across muscle activity states. **a**, Histogram of the depth below pia of recorded neurons across all three mice (366-684 single units met selection criteria). Inset is a schematic coronal section showing placement of the Neuropixel in CFA. **b**, Fractions of neurons assigned to narrow- (87-189 per mouse) and wide-waveform (279-495 per mouse) subsets in each of three mice (black circles), and the mean across animals (red bars). **c**, Normalized absolute activity change from baseline summed across the top three principal components (PCs) for recorded CFA neurons, and the top PC for muscle activity, for individual mice (thin gray) and the mean across mice (black). **d**, Scatterplot of the mean firing rate for neurons during periods of no muscle activity (rest), and periods of muscle activity, across all three mice. **e**, Fraction of neurons in different layers whose firing rate time series were significantly correlated with the activity time series for at least one forelimb muscle. One mouse had very few neurons assigned to layer 6, preventing a reliable calculation in that case. **f**, Schematic of the calculation of an activity map for each recorded neuron that is registered to inactivation effect maps. **g**, Example neural activity maps exhibiting somewhat sparse firing (black values) for 11 neurons from one mouse, along with their maximum firing rates (red). **h**, Neural maps whose sparsity values reflect quartiles of the sparsity distribution for the given mouse. **i,** Cumulative histograms of sparsity values for all neurons in each of three mice. **j**, Scatterplot of sparsity versus mean firing rate during climbing for neurons in all three mice. R2 is for a linear fit (red). **k**, Mean 2D correlation of activity maps for neurons assigned to different layers (red), compared to a null distribution computed by repeating the calculation 100 times after randomly permuting neuronal layer labels (gray), and the means thereof (black bars).

To enable alignment with CFA influence, we measured variation in the firing of neurons across the muscle activity state maps used for quantifying inactivation effects (Fig. 5f). Muscle activity state vectors for 50 ms epochs during Neuropixels recordings were embedded on the same state maps used for quantifying inactivation effects in the given animal. For each neuron, its average firing rate was estimated at each grid point, producing a “neural activity map.” This aligns neural activity during muscle activity states with inactivation effects that immediately follow similar muscle activity states.

Neural activity maps showed a wide array of muscle state-dependent firing patterns. In particular, we found many neurons with firing that was somewhat sparse across maps; firing was heavily concentrated in subregions of the maps, and mostly or completely absent in others (Fig. 5g). To quantify this sparsity, we used an index that was originally developed to measure the place-dependent firing of hippocampal neurons^54^. Ordering neurons by this sparsity index revealed that even the neurons with a median level of sparsity had firing that was heavily concentrated in subregions of the maps (Fig. 5h,i; Extended Data Fig. 6b). Sparsity was only weakly dependent on mean firing rate during climbing (Fig. 5j). Neural activity maps did not vary substantially across cortical layers, as the average 2D correlation between maps for neurons assigned to different layers was similar to that expected assuming no variation across layers (Fig. 5k). Thus, a substantial fraction of CFA neurons across cortical layers each fire primarily at a limited range of muscle activity states during climbing. These results indicate that CFA neuron firing carries information about muscle activity states organized by their similarity.

### CFA activity components that align with inactivation effects

We then used neural activity maps to identify components of CFA firing that align with CFA influence on muscles. We did so by combining singular value decomposition (SVD) and canonical correlation analysis (CCA) to align neural activity maps with the inactivation effect maps computed for the same animals^55^. The neural activity map for each wide-waveform neuron with overall mean firing rate above 0.1 Hz was converted to a vector by concatenating its columns. For each mouse, these vectors were combined into a grid points by neurons matrix, which was then replaced with a dimensionally-reduced grid points by 20 neural singular vectors matrix computed with SVD (Fig. 6a,b). These dimensionally-reduced matrices were then aligned through CCA with grid points by muscles matrices formed similarly by concatenating the columns of inactivation effect maps for the four recorded muscles (Fig. 6a-d).

**Figure 6.**
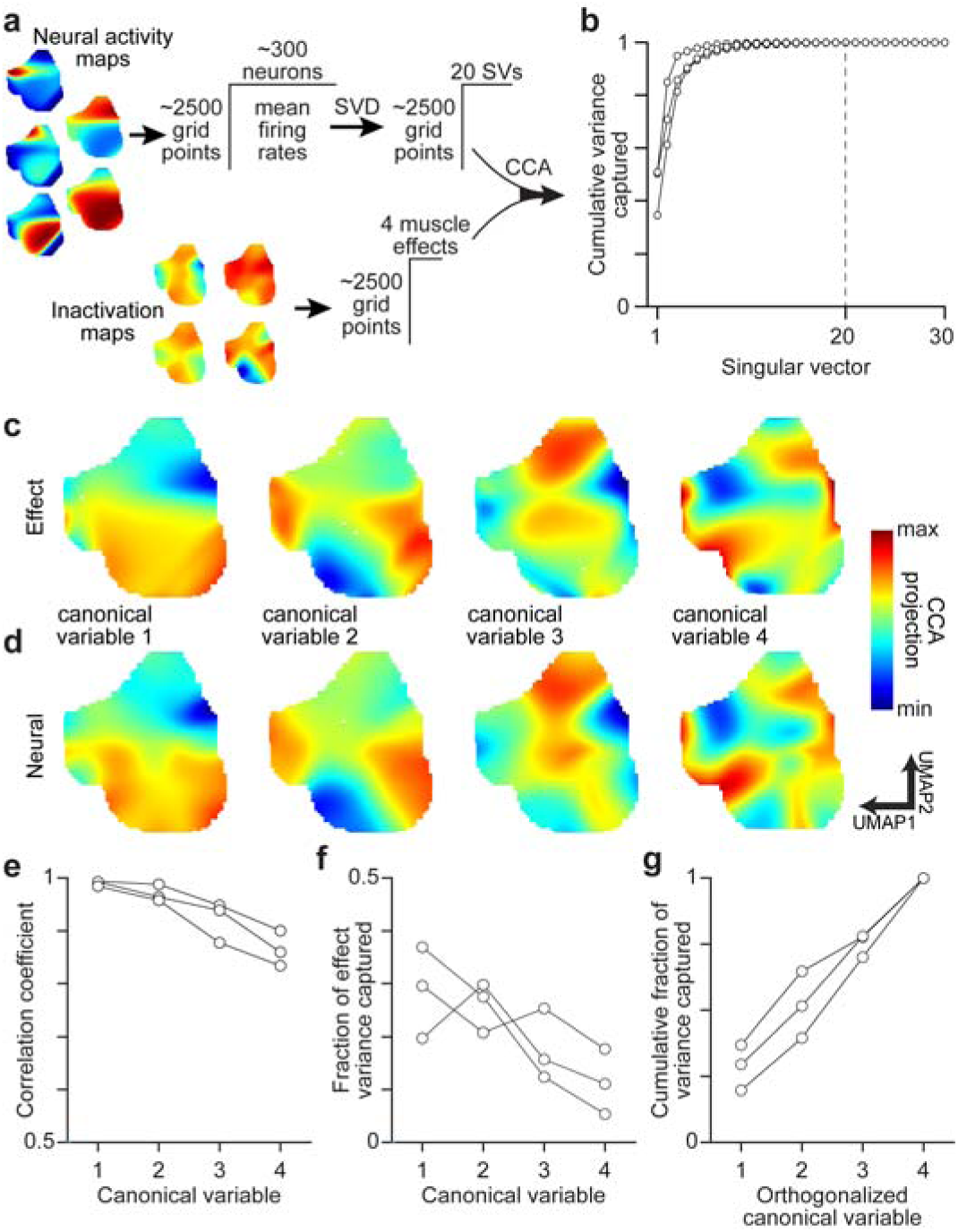
A CFA activity subspace that aligns with CFA influence. **a**, Schematic of computing the influence subspace. **b**, Cumulative neural activity variance captured for singular vectors ordered by their singular values, for each of three mice. Each connected set of dots in **b** and **e**-**g** is from a separate mouse. **c**,**d**, Canonical variables for the inactivation effects (**c**) and neural activity (**d**) for one mouse. **e**, Correlation coefficient of canonical variables for each mouse. **f**, Fractional inactivation effect variance captured by canonical variables for each mouse. **g**, Cumulative fraction of inactivation effect variance captured by canonical variables after orthogonalizing their corresponding vectors, for each mouse.

The resulting canonical variables reflect components of CFA firing patterns that maximally correlate with components of inactivation effects but are mutually uncorrelated with one another. For all three animals, neural and inactivation effect canonical variables were highly correlated, and the inactivation effect variables captured substantial fractions of inactivation effect variance (Fig. 6e,f; mean correlation = 0.99, 0.97, 0.92 and 0.86 for canonical variables 1 through 4; mean effect variance capture = 0.29, 0.26, 0.18 and 0.11). Plots of the cumulative variance captured across orthonormalized canonical vectors indicated that each inactivation effect variable captured a substantial amount of additional inactivation effect variance (Fig. 6g). Although CCA attempts to maximize correlation between canonical variables, it is not fated that each inactivation effect variable will account for a great deal of the variance in effect maps, as they do here. Repeating CCA using the inactivation effect map of each individual muscle found a CFA activity component highly correlated with the inactivation effects for the given muscle in all cases (median correlation = 0.97, range = 0.89 to 0.99, 12 total muscles in 3 animals). This indicates that CFA activity aligns well with the effects on each individual muscle.

To validate our results, we repeated CCA calculations after randomly permuting the map locations of muscle activity states from neural recordings. The resulting correlation between canonical variables was significantly lower than for the original data (for canonical variables 1 through 4, mean control correlation = 0.74, 0.72, 0.69, 0.65; p = 0.001 in all 4 cases). We also repeated CCA calculations 300 times using separate, randomly-chosen halves of trials for computing the inactivation effect maps. The canonical variables identified with each half of trials were highly similar (Extended Data Fig. 7a), as were the 4D neural activity subspaces spanned by the neural canonical vectors (Extended Data Fig. 7b). In addition, we found that CCA results were not highly dependent on the number of singular vectors used for neural dimensionality reduction (Extended Data Fig. 7c-f), nor did they depend on the choice of key inactivation map parameters (Extended Data Fig. 7g). Thus, neural canonical vectors span a neural activity subspace where activity aligns closely, and nontrivially, with inactivation effects. Below we refer to this as the “influence subspace.”

We also assessed whether subsets of neurons contributed disproportionately to these influence subspaces. To compute the relative contribution of each neuron to each canonical vector, we used the weights of each neuron in the singular vectors, and the weights of these singular vectors in the neural canonical vectors. However, we found no evidence that neurons cluster in terms of their contribution sizes (Extended Data Fig. 8a-d). Contributions to canonical vectors were substantially overlapping for neurons localized to different layers, though they were significantly higher for neurons localized to layers 5 and 6 compared to those localized to superficial layers (Extended Data Fig. 8e,f).

### Influence subspace differs from muscle activity and limb kinematic equivalents

Finally, we sought to test the hypothesis that CFA’s direct influence on muscles is mediated by the CFA activity components that correlate with muscle activity or limb kinematics in total. If this were true, we would expect that the influence subspace would be similar to subspaces aligned with muscle activity or limb kinematics (Fig. 7a). To find a subspace in which CFA activity aligns with muscle activity, average muscle activity maps (Fig. 2m) for each mouse were converted to vectors, assembled into a grid points by muscles matrix, and aligned via CCA with the matrix of dimensionally-reduced neural activity (Fig. 7b,c; Extended Data Fig. 9a,b). To find a subspace in which CFA activity aligns with limb kinematics, we used the limb kinematic state maps generated for inactivation sessions (Figure 4). We embedded vectors composed of limb site positions and their corresponding first derivatives from neural recording sessions into these existing limb kinematic state maps. We then made maps of the average horizontal and vertical positions of each site at grid points defined on the limb kinematic state maps (Extended Data Fig. 9c). We assembled neural activity maps for each neuron using segments of their firing rate time series that correspond to embedded limb kinematic state vectors (Extended Data Fig. 9d). These site position and neural maps were aligned via SVD and CCA (Fig. 7d,e; Extended Data Fig. 9e). Substantial fractions of both muscle activity and kinematics variance were captured by canonical variables that were highly correlated with their corresponding neural canonical variables (Fig. 7f-i). Thus, the resulting neural canonical vectors span neural activity subspaces where activity aligns with either muscle activity or limb kinematics.

**Figure 7.**
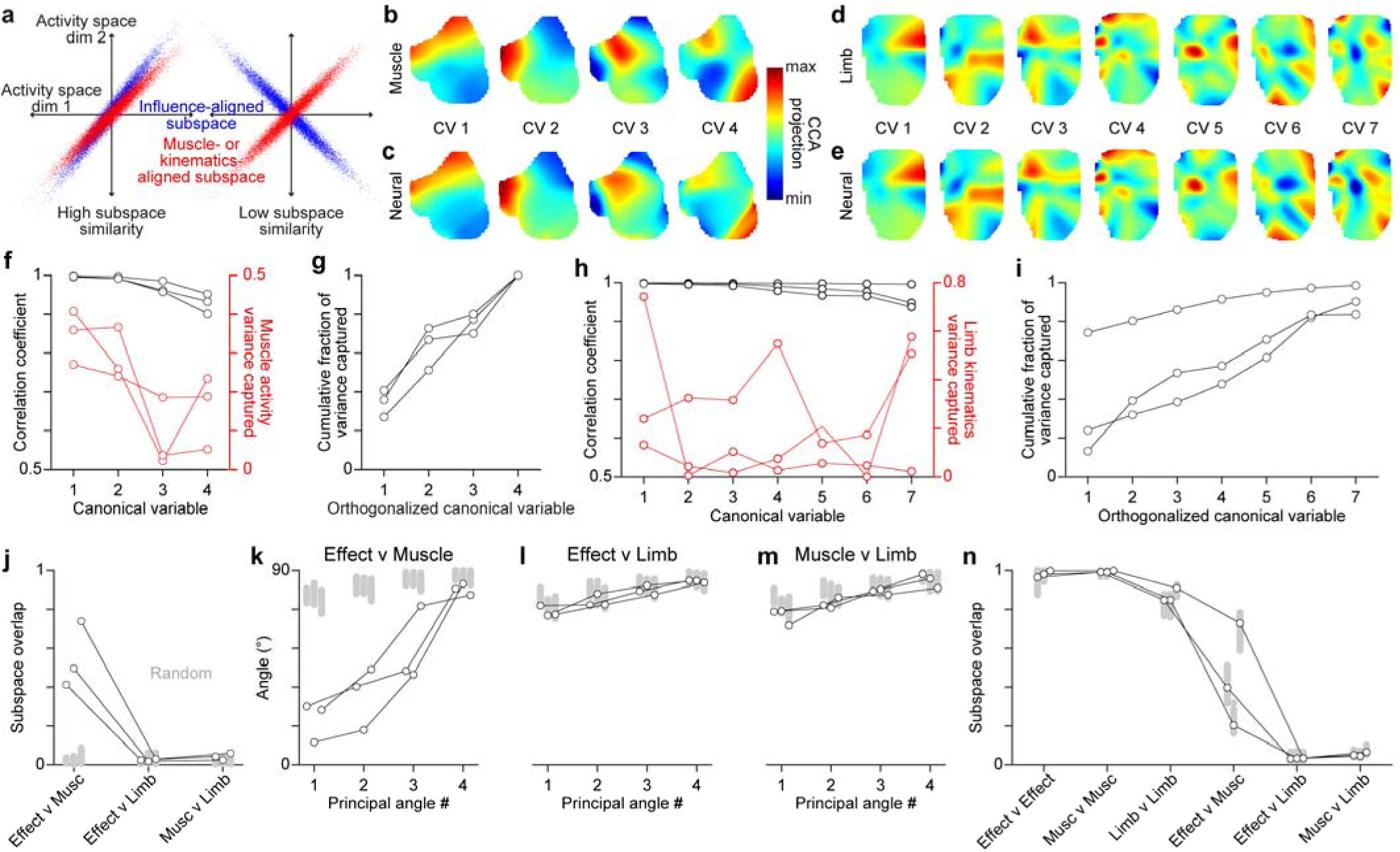
Divergence between neural activity subspaces aligned with inactivation effects, muscle activity, and limb kinematics. a, Schematic of different scenarios for the overlap between neural activity subspaces. **b**,**c**, Canonical variables for muscle activity (**b**) and neural activity (**c**) for one mouse. **d**,**e**, Canonical variables for limb kinematics (**d**) and neural activity (**e**) for the mouse used in **b**,**c**. **f**, Correlation coefficient (black) and fractional muscle activity variance captured (red) for canonical variables. Each set of connected dots in **f-n** are from one mouse. **g**, Cumulative fraction of muscle activity variance captured by canonical variables after orthogonalizing their corresponding vectors. **h**, Correlation coefficient (black) and fractional limb kinematics variance captured (red) for canonical variables. **i**, Cumulative fraction of limb kinematics variance captured by canonical variables after orthogonalizing their corresponding vectors. **j**, Overlap of different activity subspaces (black circles) compared to 300 estimates of the overlap expected by chance for each animal (gray dots). **k-m**, Principal angles for different activity subspaces (black circles) compared to 300 estimates of the principal angles expected by chance for each animal (gray dots). **n**, Mean overlap (black circles) between subspaces defined from two maps made with separate halves of time series segments, for 300 different segment parcellations (gray dots).

To evaluate the similarity between the influence, muscle activity, and limb kinematics subspaces, we compared them in two ways. First, we asked whether they were more similar than would be expected by chance for different subspaces. We measured the overlap between pairs of subspaces and compared it to the overlap when one was replaced by a random subspace that captured the same amount of neural activity variance (see Methods). On a scale from 0 (no) to 1 (complete) overlap, the overlap of the influence subspace with the muscle activity subspace ranged from 0.423 to 0.740 across the three mice, substantially above chance (Fig. 7j). However, the overlap between influence and limb kinematics subspaces ranged only from 0.018 to 0.030, and the overlap between muscle activity and limb kinematics subspaces ranged only from 0.025 to 0.060; these overlaps were on par with those expected by chance. The same relationships between subspaces were reflected in their principal angles as well (Fig. 7k-m). Overlap values were not strongly sensitive to the precise choice of key map parameters (Extended Data Fig. 9f).

Second, we asked whether these three subspaces were less similar than would be expected by chance for two subspaces of the same type. We measured the overlaps between subspaces of the same type or different types, each computed from separate sets of time series segments (Fig. 7n). In all cases, the overlap for subspaces of different types was much lower than the overlap for the same type. Thus, the influence subspace overlaps partially, but not completely, with the muscle activity subspace, yet the influence subspace has no appreciable overlap with the limb kinematics subspace.

## Discussion

Here we have assessed the direct influence of motor cortical output on muscle activity during naturalistic climbing in mice. By quantifying this influence across the full range of muscle activity states that occur during climbing, we have shown that CFA acts selectively, instructing motor output patterns. CFA activates physiological flexor muscles to varying degrees and only at a subset of muscle activity states, while activating or suppressing their antagonists less frequently. We have also shown that many CFA neurons are primarily active at subsets of similar muscle activity states. Finally, our approach enabled us to distinguish a neural activity subspace aligned with CFA influence from those aligned with muscle activity or limb kinematics. These results suggest that during an ethologically relevant motor behavior, mouse motor cortex appears to selectively direct muscle activity through a neural activity subspace distinct from those to which functional significance has previously been imputed^31–35^.

Motor cortex has activity components that are predictive of muscle activity and movement kinematics^25–28^. This could reflect a cortical role in generating all limb muscle commands during motor cortically dependent behaviors. At the same time, disturbance to motor cortical output often causes hypometric limb movements^9,24,43,56,57^, suggesting that motor cortex contributes to driving muscle activity but does not play a necessary role in determining its pattern in many cases. In either of these scenarios, we would have observed inactivation effects that were pervasive across all muscle activity states. Instead, we have found that during a cortically-dependent task in mice, the influence of primary motor cortex on forelimb muscle activity is selective and instructive. This is supported by our observations that inactivation effects on individual muscles are only present at a subset of muscle activity states, these effects vary in magnitude across that subset, and the pattern of effect magnitudes across states itself varies across muscles. To focus on CFA’s direct influence on muscles, we measured inactivation effects at the shortest latency CFA output can affect forelimb muscles in mice^18^. Effects at this latency are not seen during all motor behaviors.

Motor cortex may confer an added level of muscle control that improves movement efficacy by generating muscle activity patterns that cannot be achieved by other motor system regions^19^. Our results suggest that this role may be mediated by a selective modulation of ongoing motor output, one that differs categorically between muscles of different functional types. We were surprised to observe that direct motor cortical influence on physiological flexors was always an activation, and that influence on their respective antagonists differed not just in prevalence or magnitude, but also sign, showing a mix of activation and suppression. Given the non-stereotyped and adaptive nature of the movements mice perform in our paradigm, we had expected both activation and suppression of all muscles. For example, if motor cortex was introducing corrections to reduce the deviation from a target movement, as predicted by current theory^58,59^, we would expect both activation and suppression if errors are symmetric around the target.

It remains unclear how well our basic results here generalize to other mammals, including primates. Despite substantial homology between rodent and primate motor circuits^2,60^, functionally salient differences exist, especially in the circuits that govern finger movements^13,61^. That said, at least for non-human primates, broad swaths of the motor behavioral repertoire recover after motor cortical lesion, including walking, climbing, jumping, and even goal-oriented manual tasks, though performance efficacy is reduced in all cases^8,62,63^. These results could reflect that aside from individuated finger movements, motor cortex does not generate the entirety of limb muscle activity patterns, but instead selectively modulates muscle activity patterns to improve movement efficacy.

We found that some CFA neurons fire preferentially for different subsets of similar muscle activity states. This does not align with the view that M1 activity is well-described as a low-dimensional, linear dynamical system^34,53^. In this view, the activity of each M1 neuron reflects a linear combination of a small number of latent variables. Neurons active during distinct subsets of muscle activity states would require distinct latent variables to capture their activity, increasing the dimensionality of the population activity as a whole. A small number of linear latent variables might still capture much of the variance in our activity measurements, but they would not explain the highly state-specific firing patterns of some neurons. Because these firing patterns carry information about motor output, they may play an important role in motor control. The high state specificity of many neurons is also not predicted by previous characterizations of M1 neurons as broadly tuned for muscle activity or movement kinematics^28,64,65^. Here again, though, this aspect of our results may not generalize to other species, or to the ballistic reaching tasks used in many previous studies. That said, it does align with recent demonstrations of reproducible high-dimensional activity in monkey M1^66^.

Our observations also have important implications for the functional interpretation of kinematics-related neural activity. By focusing on a context in which muscle activity and gross limb kinematics are substantially uncoupled, we found that the neural activity subspace where activity aligns with these kinematics is essentially orthogonal to the one that aligns with direct CFA influence. However, we cannot rule out that there are other, perhaps finer, kinematic features that more substantially covary with direct CFA influence. Despite this, our observations remain a significant revelation given the contemporary prevalence of video-based kinematic tracking for measuring nervous system output. Prominent kinematic features can correlate substantially with activity in a neuronal population, but correlate negligibly with the functional influence of that activity.

## Online Methods

All experiments and procedures were performed according to NIH guidelines and approved by the Institutional Animal Care and Use Committee of Northwestern University.

### Experimental Animals

A total of 50 adult male mice were used, including those in early experimental stages to establish methodology. Strain details and number of animals in each group are as follows: 44 VGAT-ChR2-EYFP line 8 mice (B6.Cg-Tg(Slc32a1-COP4*H134R/EYFP) 8Gfng/J; Jackson Laboratories stock #014548); and 6 C57BL/6J mice (Jackson Laboratories stock #000664).

All mice used in experiments were individually housed under a 12-hour light/dark cycle in a temperature- and humidity-controlled room with *ad libitum* access to food and water, except during experiments. At the time of the measurements reported, animals were 12-18 weeks old. Animals weighed 24–30 g. All animals were being used in scientific experiments for the first time. This included no previous exposure to pharmacological substances or altered diets.

### Climbing apparatus

The climbing apparatus (Extended Data Fig. 1) was housed inside a sound attenuating chamber (H10-24A, Coulbourn). Experimental control was performed using the Matlab Data Acquisition Toolbox, the NI PCI-e-6323 DAQ, and two Arduino Duos. The climbing apparatus itself consisted of a 3D-printed cylindrical wheel with alternating handholds positioned 12 degrees apart from each other. The right handholds were affixed to linear actuators (L-12-30-50-12-I, Actuonix) while the left handholds were statically positioned. A ratchet mechanism was used to ensure that the climbing wheel could only rotate downwards from the mouse. One end of the wheel was supported by a shaft angular encoder (A2-A-B-D-M-D, U.S. Digital). Angular position signals were sent to the Arduinos to track the location of each handhold. When each right handhold reached a position 180° away from the mouse, the linear actuator moved the handhold to a new, randomly chosen mediolateral position. The other end of the wheel was supported by a slip ring (SR20M-LT, Michigan Scientific) that commuted voltage signals to and from the actuators embedded in the wheel. Water rewards were dispensed with a solenoid valve (161T012, NResearch) attached to a lick tube (01-290-12, Fisher), and this dispensation was controlled by Matlab through the NI PCI-e-6323 DAQ. A speaker was used to play a 5 kHz tone for 200 ms whenever rewards were dispensed.

### Training

Under anesthesia induced with isoflurane (1–3%), mice were outfitted with titanium or plastic head plates affixed to the skull using dental cement (Metabond, Parkell). Headplates had an open center that enabled subsequent access to the skull, which was covered with dental cement. During headplate implantation, the position of bregma relative to marks on either side of the headplate was measured to facilitate the positioning of craniotomies during later surgeries. After recovery from headplate implantation surgery, mice were placed on a water schedule in which they received 1 ml of water per day.

At least 4 days after the start of the water schedule, mice were acclimated to handling by the experimenter following established procedures^67^. After acclimation to handling, mice were acclimated to head-fixation over two daily sessions during which they were placed in a 3D printed hutch positioned directly in front of the climbing wheel apparatus and provided water rewards (3 μl per reward) at regular intervals.

Following acclimation, mice underwent daily hour-long training sessions on the wheel apparatus. Training involved an initial stage (1-3 sessions) aimed at training mice to grab for and pull at the handholds in order to rotate the wheel downward and receive water reward. Mice were head-fixed in an upright position facing the front of the wheel so that all four limbs could easily grab onto the handholds, and the right handholds remained fixed. Rewards were triggered by an experimenter’s key press whenever a mouse performed any slight rotation of the wheel downward towards his body, and longer or faster bouts were rewarded with additional rewards. Over the course of these sessions, the mice generally learned to associate rotating the wheel with a water reward and began iteratively rotating the wheel.

During the next stage of training (4-10 sessions, median 5), right handholds were kept fixed and mice were encouraged to rotate the wheel for increasingly long bouts. Here, rewards were dispensed for continuous climbing bouts above a threshold distance after the bout ended, using an automated experimental control script written in Matlab. The first ten times during a training session that the threshold distance was met, mice automatically received a water reward. Subsequently, the total distance traveled was compared to those from the previous 10 bouts. If the time was above the 25th percentile value for those 10 bouts, the mouse received one water reward. If it was above the 60th percentile value, the mouse received two water rewards. And if it was above the 90th percentile value, the mouse received four water rewards. Otherwise, the mouse received no water reward. The threshold distance was adaptively adjusted to maintain the reward rate such that the mouse received approximately 1 mL of water during each hour-long training session. Thus if the recent reward rate was too low, the threshold distance was lowered, and if the recent reward rate was too high, the threshold distance was raised. During all subsequent training sessions, the right handhold positions were randomly repositioned along the horizontal axis after rotating past the mouse, though the same reward scheme was used.

In this paradigm, mice perform a sophisticated locomotion that deviates from rhythmic pattern to negotiate unpredictable variation in the “terrain” presented by the handholds. The consistency of climbing form and performance across sessions (Figure 1), together with the natural climbing ability mice possess, obviate the need for extensive training. In fact, some mice exhibit long, continuous bouts of climbing during the first session during which they are taught the climbing-reward pairing.

### Electromyographic recording

Electromyographic (EMG) electrode sets were fabricated for forelimb muscle recording using established procedures^18,39^. Briefly, each set consisted of four pairs of electrodes. Each pair was comprised of two 0.001″ braided steel wires (793200, A-M Systems) knotted together. On one wire of each pair, insulation was removed from 1 to 1.5 mm away from the knot; on the other, insulation was removed from 2 to 2.5 mm away from the knot. The ends of the wires on the opposite side of the knot were soldered to an 8-pin miniature connector (11P3828, Newark). Different lengths of wire were left between the knot and the connector depending on the muscle a given pair of electrodes would be implanted within: 3.5 cm for biceps and triceps, 4.5 cm for extensor carpi radialis and palmaris longus. The ends of wires with bared regions had their tips stripped of insulation, then were twisted together and crimped inside of a 27-gauge needle that facilitated insertion into muscle.

Mice were implanted with EMG electrodes during the surgery in which headplates were attached. The neck and right forelimb of the mouse were shaved, and incisions were made above the muscle to be implanted. Electrode pairs were led under the skin from the incision on the scalp to the incision at the site of implantation. Using the needle, electrodes were inserted through the muscle, and the distal portion of the electrodes was knotted to secure both 0.5 mm bared wire regions within the muscle. The needle and excess wire were then cut away. Incisions were sutured and the connector was affixed with dental cement to the posterior edge of the headplate.

EMG recordings were amplified and digitized using a 16-channel bipolar amplifying headstage (C3313, Intan Technologies). Data was acquired at 4 kHz using the RHD2000 USB interface board (Intan Technologies).

### Optogenetic inactivation

After a VGAT-ChR2-EYFP mouse completed a few climbing sessions with randomly positioned handholds, dental cement above the skull was removed and a 2-mm-diameter craniotomy was made above the left caudal forelimb area centered at 1.5 mm lateral and 0.25 mm rostral of bregma. A thin layer of Kwik-Sil (World Precision Instruments) was applied over the dura and a 3-mm-diameter #1 thickness cover glass (Warner Instruments) was placed on the Kwik-Sil before it cured. The gap between the skull and the cover glass was then sealed with dental cement around the circumference of the glass. A custom stainless steel ferrule guide (Ziggy’s Tubes and Wires) was then cemented to the headplate a certain distance above the surface of the brain. This distance was set to ensure that the cone of light emanating from a 400 μm core, 0.50 NA optical patch cable terminating in a 2.5 mm ceramic ferrule (M128L01, Thorlabs) would project a spot of light 2 mm in diameter onto the surface of the brain (Fig. 2a). The ferrule guide enabled quick and reliable positioning of the ferrule above the brain surface so that a large expanse of cortex could be illuminated. In previous experiments using this method for inactivation, control experiments in wildtype mice showed no discernible muscle activity perturbation in response to light^18^. Moreover, the short latency at which we measure effects here would preclude visually-driven responses.

To attenuate firing throughout motor cortical layers, we used a 450 nm laser (MDL-III- 450, Opto Engine) to sporadically apply 25 ms light pulses at an intensity of 10 mW/mm^2^ to the brain surface. During each session that involved inactivation of the motor cortex, trials were initiated after the distance climbed on the current bout exceeded a random threshold between 0° and 20°, and a mouse was actively climbing. Light was applied on a random third of trials. This ensured that inactivation and control (no light) trials were broadly distributed across the muscle activity states that occur during climbing. Inactivation trials were never triggered less than 5 seconds apart. A total of 2292-5115 trials (median = 2715) were collected in each mouse (roughly 1/3 inactivation and 2/3 control), spanning 11-37 (median = 18.5) climbing sessions. Including twice as many control trials as inactivation trials enabled comparisons between separate sets of control trials to aid statistical testing.

### Neural recording

For some mice, optogenetic inactivation sessions were followed by 3-4 daily neural recording sessions that typically lasted an hour each. One day prior to the first neural recording session, the cover glass and Kwik-Sil were removed, a small durotomy was made in the craniotomy center, and a Pt/Ir reference wire was implanted to a depth of 1.5 mm in left posterior parietal cortex. Opaque silicone elastomer (Kwik-Cast, World Precision Instruments) was used to cover the craniotomy after surgery and between recording sessions. At the time of recording, the exposed brain surface was covered with agarose and silicone oil or liquid paraffin oil. A Neuropixel 1.0 (IMEC) was subsequently inserted to a depth of 1.5 mm into the brain (Fig. 5a) at a rate of 3-5 μm/s using a motorized micromanipulator (MP-225A, Sutter Instrument). Electrode voltages were acquired at 30 kHz and bandpass filtered at 0.3 to 10 kHz using SpikeGLX (Bill Karsh, https://github.com/billkarsh/SpikeGLX), and then sorted with Kilosort 2.0^68^ (https://github.com/MouseLand/Kilosort2). Sorted units were assigned to separate cortical layers based on the depth below pia assigned to the waveform centroid in Kilosort and the laminar depths reported in^69^. To be conservative, units near laminar boundaries were ignored in analyzing neural responses by layer. To classify units as either wide-waveform, putative pyramidal neurons or narrow-waveform, putative interneurons, we pooled the spike widths of all sorted single units, obtaining a bimodal distribution^18,70^. Neurons with spike widths > 450 µs were classified as putative pyramidal neurons, and the remainder were classified as putative interneurons. The proportion of units classified as wide- and narrow-waveform was in line with previously observed proportions^13,44^

### EMG preprocessing

For both optogenetic inactivation and Neuropixel recording sessions, EMG measurements were downsampled to 1 kHz, high-pass filtered at 250 Hz, rectified, and convolved with a modified Gaussian filter kernel. We used causal filtering to enable precise assessment of perturbation latencies, and hence used a Gaussian filter kernel that was initially defined with a 10 ms standard deviation, but then had amplitudes for times < 0 – its “backwards in time” side – set uniformly to zero. EMG traces were then z-scored across all time points during the given session.

### Video recording and analysis

A high-speed, high-resolution monochrome camera (Blackfly S USB 3.0, 1.6 MP, 226 FPS, Sony IMX273 Mono; Teledyne FLIR, Wilsonville, OR) with a 12 mm fixed focal length lens (C-Mount, Edmund Optics) was positioned to the right of the head-fixed mouse during inactivation and neural recording sessions, and videos were acquired under a near-infrared light source at 100 frames per second with a resolution of 400 by 400 pixels. During optogenetic inactivation sessions, the camera was triggered to start recording using StreamPix software (NorPix, Inc; Montreal, Quebec, Canada) 20 ms before each inactivation or control trial. Each recording lasted 500 ms beyond the 25 ms light or control command pulse that marked the trial. Annotation of behavior was accomplished using DeepLabCut^71^. To enable better markerless tracking, the right forelimbs of mice were shaved and tattooed (Black tattoo paste, Ketchum Mfg., Lake Luzerne, New York) at 8 different sites along the right arm. All videos were also adjusted with ffmpeg (ffmpeg.org) to improve brightness and contrast. DeepLabCut using the ResNet-50 neural network with an Adam optimizer was trained on the annotated images for 1,030,000 iterations; on ∼4000 randomly sampled video frames across mice and sessions, we provided manually labeled locations of the 8 forelimb sites for training: shoulder, two sites between the shoulder and elbow, elbow, two sites between the elbow and wrist, wrist, and tip of the last digit (Fig. 4b). The training set comprised 80% of the labeled frames.

All DeepLabCut-tracked forelimb site trajectories were then exported to Matlab for further post-processing to remove outliers (Extended Data Fig. 10). First, sites in each video time series that were assigned by DeepLabCut a likelihood (i.e. its confidence that a site was correctly labeled) lower than 0.75 were replaced with an interpolated value using the median of the ten previous and ten following values (Matlab function fillmissing). Next, outliers in site position time series were identified using the median absolute deviation (MAD): shoulder coordinates were constrained to lie within 1.5 MAD from their median, digit coordinates to be within 3 MAD from their median, and all other joints to be within 2 MAD from their respective medians. Outliers were replaced with an interpolated value using a moving median of window length 10. Lastly, limits were imposed on the pairwise distances and angles between neighboring joints (the shoulder-elbow, the elbow-wrist, and the wrist-digit tip) such that the angle between shoulder-elbow and elbow-wrist could not exceed 180 degrees, and the distances between each of these joints were within 2 MAD of their medians. Site positions not meeting these criteria was also replaced with an interpolated value using a moving median of window length 10.

We note that our findings here apply only to the kinematic features our measurements have captured, primarily the right forelimb joint angles in the sagittal plane. Our measurements do not account for forelimb orientation in the mediolateral dimension. However, the elbow and wrist muscles we have recorded from primarily govern the angles of the elbow and wrist, which our measurements do capture. Mediolateral forelimb orientation is primarily determined by shoulder muscles. Moreover, as illustrated in the Supplementary Movie, the range of limb motion in the two dimensions we did examine is much larger than the range in the mediolateral dimension. Thus, inclusion of the mediolateral dimension in measurements is not likely to have changed our results substantially.

### Behavioral analysis

For each animal, we computed the principal angles between muscle activity time series for each of the first 20 climbing sessions that used the automated control script, compared to the 20^th^ session. PCA was first performed on the muscle activity time series for the four recorded muscles. The top 2 PCs were used for the subsequent calculations, since they captured over 90% of activity variance. Principal angles were then computed using a standard approach: The cross-covariance matrix was computed for the two PC time series from the given session and those from the 20^th^ session, and SVD was applied to the result. The principal angles were then computed as the real part of the inverse cosine of the resulting singular values.

Sample entropy^41^ was computed for the muscle activity time series for the four recorded muscles using the ‘sampen’ function downloaded from the Matlab file exchange, with the embedding dimension set to 10. Sample entropy has previously been used to measure regularity in signals like ECG, EEG, and BOLD from fMRI.

### Inactivation effects during stereotypical climbing features

We identified three recurring kinematic features that are stereotypical in our observations of climbing behavior – pulling a handhold down with the right forelimb, reaching the right forelimb up for a handhold, and palpation with the right hand when grasping for a handhold. Limb movement during each of these features was best reflected along the Y axis of video frames. Pulling down was accompanied by an increase in averaged Y coordinates as measured in pixels (values increase as you move downward in the video frame), which corresponds to a positive slope. The opposite was true for reaching up. Handhold palpations were associated with an oscillatory increase, decrease, and increase, or vice versa, of Y coordinates; both of these patterns correspond to a minimum of two sign changes in slopes over a brief time window. To identify instances of each feature during climbing, we thus looked at changes in the Y coordinates of the four tracked forelimb joints (shoulder, elbow, wrist, and finger) across video recordings spanning 100 ms before and after trial onset. We first fit a line of best fit to the Y coordinates averaged over the four tracked joints across the entire 200 ms window to obtain a slope. Then, we calculated slopes from overlapping 50 ms sliding windows that began every 10 ms, starting from 100 ms before trial onset. From these slope measurements, we detected changes in the slope across time. A trial was assigned as pulling down if the overall slope was positive with a magnitude of 2 or greater and no sign changes in slope were detected. Conversely, a segment was assigned as reaching up if the overall slope was negative with a magnitude of −2 or less and no sign changes were detected. Segments with three or more slope sign changes were assigned as handhold palpation. Following this, for verification purposes, we randomly selected a quarter of trials assigned to each of the three features. Using visual inspection of video data, we confirmed that the kinematics associated with each trial matched the assigned feature in all cases. This validated the criteria we imposed for assignment. Trial averages of the muscle activity and limb kinematic time series aligned by trial onset were then assembled for each behavioral feature.

### Muscle activity state maps

We explored organizing inactivation and control trials based on the forelimb muscle activity immediately preceding trial onset. For example, we plotted trials along axes defined by the relative activity of antagonist muscles at each joint (Extended Data Fig. 3d) or along the top two principal components for the activity of all four muscles (Extended Data Fig. 3e). However, the density of trials across these plots varied greatly, with subsets of trials clustered closely together. Thus these plots were not conducive to collecting together trials in similarly sized groups that would afford a similar degree of statistical power; dividing trials based on these plots would concentrate trials, and statistical power, around certain states. To more effectively distribute the statistical power afforded by our trials, we explored the use of dimensionality reduction methods that organize states according to their *n* nearest neighbors.

We ultimately used UMAP to obtain 2D muscle activity state maps on which nearby states are highly similar. We applied UMAP to segments extracted from the EMG time series collected during the optogenetic inactivation sessions, and their corresponding first derivatives. For each control or inactivation trial, we defined 13 overlapping 50 ms epochs centered every 10 ms from −55 to +75 ms from command pulse onset (Fig. 2c). Then, for each 50 ms epoch, we averaged the EMG traces for each muscle over 5 ms bins, and concatenated the resulting values for the four muscles and their corresponding first derivatives, yielding one 80 by 1 vector. Thus in these vectors, the first 40 values reflect the EMG signals from 4 muscles, and the last 40 values reflect their first derivatives (Fig. 2c). UMAP (Matlab function run_umap, from the Matlab File Exchange, with the following parameters: n_neighbors = 30, n_components = 2, min_dist = 0.3, metric = Euclidean) was applied to embed all resulting vectors from both inactivation and control trials, generating the 2D muscle activity state maps (Fig. 2d). Using two dimensions here simplified subsequent quantification of inactivation effects across muscle activity states. Using overlapping epochs ensured that embedded state vectors from individual trials formed continuous trajectories across the resulting maps (Fig. 2d).

### Inactivation effect maps

To quantify the influence of CFA inactivation on muscles across the muscle activity state maps, we first excluded outlying embedded state vectors. For each embedded vector on a given map, we computed its mean Euclidean distance on the map from all other vectors. A vector was deemed an outlier if its mean distance was more than three standard deviations above the mean for all other vectors. Some mice had no embedded vectors classified as outliers, and the percentage classified as such was overall less than 1%.

We then defined uniformly spaced grid points across each map at which we would calculate the effects of CFA inactivation. Since the number of grid points of a fixed spacing across a map would depend on the scale of the map, which could vary between maps, we rescaled the coordinates of each 2D map rd to 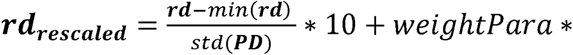2, where *PD* is the set of all Euclidean distances between embedded vectors, and weightPara is the standard deviation of the Gaussian function used to weight trials in computing the inactivation effect size at each grid point (see below; “spatial filter width” in Extended Data Figs. 3, 6 and 8). For all maps, we used weightPara = 5.

We then sought to compute the inactivation effect at each grid point, based on trials that began from muscle activity states close to each given grid point. Our approach was to compute effects for each grid point from inactivation and control trial averages made from all trials of a given type, but where trials were weighted by the distance on the 2D map from the given grid point to the weight epoch state vector for the given trial. The weight epoch state vector was defined as the embedded vector from the epoch spanning −40 to +10 ms from command pulse onset on the given trial (i.e., the epoch just before any direct effect could begin; Fig. 2c,e). However, the density of these vectors varied across maps, meaning that the summed weight of trials contributing to trial averages would vary across grid points as well. Given this, to ignore grid points around which the weight epoch state vectors were too sparse for reliable trial averaging, a grid point was designated as “valid” only if ∑_i_ W_g,i_ > a threshold. W_g,i_ was in essence a Gaussian function, defined as 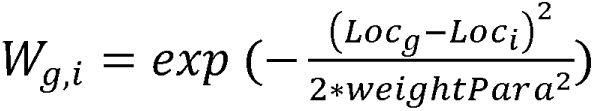, where i is the weight epoch state index (one per trial), g is the grid point index, and Loc_g_ - Loc_i_ is the Euclidean distance between the weight epoch state and the grid point on the 2D map. We used a threshold of 10, which led nearly all grid points falling within a convex hull surrounding embedded vectors to be classified as valid, while nearly all grid points falling outside this hull were not. Grid points not designated as valid were ignored in subsequent calculations. All embedded muscle activity state vectors (those not deemed outliers) were close to valid grid points, and so this criteria did not prevent an appreciable contribution from any vectors. Together with the very small fraction of embedded vectors ignored as outliers, this implies that our analysis involves practically all the muscle activity states that occur during climbing.

Next, we calculated the trial-averaged activity for each muscle at each valid grid point, separately for the inactivation and control trials. For each muscle and trial type, we extracted segments of their activity time series from −10 to +30 ms relative to command pulse onset. We then took a weighted average of these segments across each trial type, where each segment was weighted by W_g,i_. At each grid point, this produced control and inactivation trial averages for each muscle, where each trial is weighted by the distance from the grid point to the muscle activity state just prior to any direct inactivation effect during the trial. Using weightPara = 5 here (roughly 10% of the width of the map) reflects a trade-off between differentiating effects across distinct map regions and the statistical power gained from combining trials.

Finally, we quantified the size of the CFA inactivation effect at each valid grid point by comparing the rates of change (slopes) in inactivation and control trial averages 0 to +20 ms from command pulse onset (Fig. 2f). Muscle activity at time 0 ms was defined as the mean from −10 to +10 ms, and likewise activity at +20 ms was defined as the mean from +10 to +30 ms.

Quantifying the inactivation effect by taking the ratio or difference between the control and inactivation slopes returned qualitatively similar results. We used the difference because it was more easily interpretable (Fig. 2h).

We also sought to assess the similarity between nearby state vectors embedded in local neighborhoods across maps, and thus the similarity of states from which trials contributing strongly to given grid point trial averages began. To do this, we computed the average Euclidean distance in the full 80D space between state vectors embedded near each grid point on the 2D maps. We first calculated the Euclidean distances between all possible pairs of embedded vectors in the full 80D space. For each grid point, we then computed a locally weighted average of these distances. In these averages, each distance *d* was weighted by a Gaussian function of the distance on the 2D map between the grid point and the midpoint between the corresponding two embedded vectors (i.e., those separated by *d*), using the same Gaussian function as above (weightPara = 5).

There are a number of important caveats in evaluating inactivation effect maps. Collapsing 80D muscle activity state vectors onto a 2D map eliminates substantial information about those states. The smoothing we have applied in computing trial averages will also mask fine structure. UMAP organizes states according to similarity and so ignores other features potentially relevant to cortical influence. As a consequence, these maps do not reflect all the structure that may exist in the relation between muscle activity state and CFA influence.

### Analysis of inactivation effect maps

To determine which muscle activity states were significantly influenced by CFA inactivation, we performed a two-tailed nonparametric permutation test at each grid point by computing the probability of obtaining the observed inactivation effect size by chance. For each animal, and each grid point, 300 permutations were performed by first randomly splitting control trials into two groups, each with a number of trials equal to the number of inactivation trials. The number of trials in both the control (N_eontrol_) or inactivation (N_inaetivation_) trials groups was such that if N_total_control_/2 > N_total_inaetivations_, we would set N_control_ = N_total_inaetivations_; otherwise, N_inaetivation_ = N_total_eontrol_/2. Since our experiments were designed to collect twice as many control trials as inactivation trials, control trials could be sampled without replacement during the splitting process. For each grid point, and for each permutation, we calculated the inactivation effect size using the control trial average, computed as above, for one randomly selected group of control trials and the inactivation trial average, also computed as above. For each permutation, and at each grid point, we also calculated the effect size expected by chance (“null”) using trial averages for the two control trial groups. Then, at each valid grid point, we randomly picked one inactivation effect size from the 300 permutations and compared that with all 300 null effect sizes. To calculate the p-value, we compared the 300 null effects with the randomly chosen inactivation effect, calculated the fractions of the null effects where the null effects were smaller than or greater than the inactivation effect, and multiplied the smaller fraction by 2. To correct for multiple comparisons, the Benjamini-Hochberg method was used to control the false discovery rate (Matlab function fdr_bh). The effect size at each grid point was considered to be statistically significant if the FDR-corrected p-value was less than 0.05 (i.e., the likelihood of false discovery was 5% or less). We also used p-values across all grid points to estimate the fraction of null hypotheses that were false for each mouse (Matlab function mafdr), and used this as the fraction of grid points exhibiting inactivation effects.

The 2D autocorrelation for inactivation effect maps was computed using the Matlab function xcorr2, and normalized by the maximum value for the given map. For controls, maps were generated using null effect sizes at each grid point from one of the 300 comparisons of effect sizes computed with two separate sets of control trials described above. To test that inactivation effect maps differ across muscles for each animal using the Kruskal-Wallis test, we first removed inactivation effect sizes from the maps until the minimum distance between the grid points for remaining effect sizes was greater than 3x the standard deviation of the Gaussian function used for computing grid point trial averages. This ensured that remaining effect sizes were effectively computed from separate trials. Roughly ∼15-20 inactivation effect sizes remained after this removal. Linear models were fit to scatter plots using the Matlab function fitlm. The 2D correlation between inactivation effect maps was computed using the Matlab function corr.

### Maps of average muscle activity

For each muscle, we first calculated its mean activity during the 50 ms epoch reflected in each state vector. Then for each grid point, we calculated a weighted average of these values, where the mean for each epoch was weighted by a Gaussian function of the Euclidean distance on the 2D map between the epoch’s embedded muscle activity state vector and the given grid point. Here we set the standard deviation of the Gaussian function to be the same as that used for the inactivation effect maps. The resulting plots clarify the activity of muscles at each grid point and how this activity varies across grid points.

### Neural activity maps

To compare firing patterns in CFA to inactivation effects, we generated maps of average firing rate across grid points for each recorded CFA neuron. Since neural recordings and optogenetic inactivations were carried out in separate sessions, the first step was to align muscle activity states from the neural recording sessions with those from the optogenetic inactivation sessions by embedding the former set of states on the same 2D map used to make inactivation effect maps. To do this, we identified a large number of 50 ms epochs during active climbing from the neural recording sessions. To identify these epochs, peaks in the activity summed across all four muscles were first detected using the Matlab function findpeaks, where a peak was identified if a sample in the time series surpassed a threshold equal to the mean + ½*std. Based on visual inspection, this identified many peaks that always fell during active climbing. Next, two time points (“onset points”) were randomly selected on either side of the peak, such that the expected interval from the peak to each point was 50 ms. Next, any onset point was eliminated if it was less than 50 ms from either the previous or subsequent onset point. This ensured that 50 ms epochs defined around onset points would not overlap. For each remaining onset point, EMG time series segments from the epoch spanning −45 to +5 ms relative to the onset point were extracted. Each recording session typically yielded upwards of 30,000 epochs. Of these, 10,000 were randomly selected for each session to limit compute time for subsequent calculations. State vectors (80 x 1) were assembled for each epoch as above.

We then embedded the resulting 10,000 state vectors from each neural recording session onto the muscle activity state maps previously defined via UMAP to make inactivation effect maps for the given mouse. We next extracted the segments of each neuron’s firing rate time series corresponding to each state vector. For each neuron and at each valid grid point, the firing rate segments were averaged as above, weighting each segment by a Gaussian function of the distance on the map from their corresponding state vector to the grid point (weightPara = 5). The resulting segment-averaged firing rates were then averaged across time, yielding a single scalar firing rate value for each neuron at each grid point (Fig. 5f, right, and g).

For analysis described below, only putative pyramidal neurons were included. To estimate the number of neurons that showed behaviorally-dependent firing across muscle activity states, we first generated an empirical null distribution for the degree of variation across neural activity maps separately for each neuron. To do so, we reassigned each state vector from neural recording sessions to the location of a different, randomly selected vector on the map and recomputed the neural activity maps. We repeated this 500 times to yield 500 permuted maps for each neuron. To assess behaviorally-dependent variation, the skewness of the original neural activity map values and those on the 500 permuted neural activity maps was calculated (Matlab function kurtosis) as 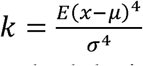, where is the set of firing rates at all grid points, is the mean of x, er is the standard deviation of x, and E(·) represents the expected value. Significance was assessed using a p-value, defined as the fraction of null distribution skewness values greater than the original. From the p-values for all neurons in a given mouse, we estimated the fraction of false null hypotheses (Matlab function mafdr) and used this as the fraction of cells with behaviorally-dependent firing.

The sparsity for neural activity maps was computed using the formula given in ^54^. Here the sparsity is the mean of the firing rates at each grid point on the map, squared and divided by the mean of the squares of the firing rates at each grid point. Note that as in the original reference, this minimizes the dependence of sparsity values on the frequency that each state on the map is visited. The 2D correlation between neural maps was computed using the Matlab function corr.

### Neural subspace for inactivation effects

Separately for each animal (n = 3 mice), neural subspaces were identified using singular vector canonical correlation analysis (SVCCA^55^) applied to the inactivation effect and neural activity maps. A singular value decomposition (SVD)-based approach was taken here because the number of recorded neurons was much larger than the number of recorded muscles. Neurons with mean firing rates lower than 0.1 Hz over the 10,000 epochs were excluded. Two matrices were generated for alignment with SVCCA. One matrix had a column for each neuron generated by vertically concatenating the successive columns of that neuron’s neural activity map. The resulting matrix had dimensions of N_grid_ by N_neuron_, where N_neuron_ is the number of neurons recorded in given mouse across recording sessions (putative pyramidal, mean firing rate over 0.1 Hz), while N_grid_ is the number of valid grid points. The second matrix was made in similar fashion by vertically concatenating the successive columns of the inactivation effect map for each muscle. This matrix had dimensions of N_grid_ by 4 since there were four muscles recorded.

SVCCA was conducted in two steps. Neural activities were first soft normalized^72^ using a soft normalization constant of 5 Hz. SVD was performed using numpy.linalg.svd in python to decompose the neural data into left singular vectors, a diagonal matrix containing singular values and right singular vectors. Next, the diagonal matrix, truncated to just the top 20 singular values, was multiplied with the corresponding top 20 right singular vectors, resulting in 20 neural activity components. CCA was applied to the N_grid_ by 20 matrix of these components and the N_grid_ by 4 matrix of inactivation effects. Twenty dimensions were retained here because the amount of variance captured and CCA alignment quality (canonical variable correlation) saturated at around 20 dimensions. CCA was also repeated with individual columns of the matrix of inactivation effect sizes in place of the full 4D matrix to show that there were CFA activity components correlated with the inactivation effects for each individual muscle.

To compute the additional variance captured by each successive canonical vector, canonical vectors were orthogonalized with the Gram-Schmidt process. In addition, we randomly shuffled the neuronal firing rate segments relative to the embedded vectors on the maps to generate shuffled neural activity maps, and performed SVCCA again as a negative control. The highest correlations between the canonical neural and effect vectors for all animals were less than 0.75. To further verify the effectiveness of SVCCA, separately for each mouse we split the inactivation and control trials from inactivation experiments randomly into two groups, calculated separate inactivation effect maps for each group, used SVCCA to find subspaces where CFA activity aligns with each of them, and calculated the principal angles between the two resulting neural activity subspaces. This procedure was repeated 300 times (Extended Data Fig. 7b).

Using the CCA results for individual columns of the matrix of inactivation effect sizes (i.e., results for effects on individual muscles), we computed the effective weight of each neuron’s activity in each canonical variable by matrix multiplying the neuron to singular vector coefficients and the singular vector to canonical vector coefficients. The four individual muscle effect size vectors yielded four effective weights for each neuron. To compare across animals in which we had recorded different numbers of neurons, we normalized these weights to have a median of 1 for each animal (Extended Data Fig. 8). To measure the relative contribution of neurons across all four muscles, we computed the norm of the 4-element vector composed of the weights for each muscle for a given neuron, and again normalized these so the median for each animal was 1.

### Overlap between subspaces

To find a neural activity subspace that aligned with muscle activity itself, we again performed SVCCA using the N_grid_ by N_neuron_ matrix of mean firing rates. However, in place of the matrix of inactivation effect sizes, an N_grid_ by 4 matrix was used where each column reflected the average activity at each grid point for one of the four muscles, computed just as was done for the neural activity maps, except muscle activity from inactivation sessions was used. Similar methods were used to find a neural activity subspace that aligned with limb kinematics. Here, instead of 4 columns, we began with 16 – the horizontal and vertical coordinates for the 8 positions tracked along the right forelimb. Since the 16 kinematic variables were highly correlated, SVD was also used on this matrix to reduce its dimensionality to 7 before performing CCA.

To compute principal angles between two neural activity subspaces, we orthonormalized the neural canonical vectors defining each subspace, computed the cross-covariance matrix for the two sets of vectors, computed the singular value decomposition of the matrix, and calculated the inverse cosine of the singular values in degrees. We also measured the degree of overlap between neural activity subspaces using the metric 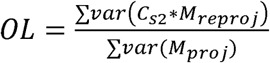, where M is the original matrix of neural activity, C_s1_ and C_s2_ are matrices comprising coefficient vectors that that define the two subspaces, M_proj_ = C_s1_ * M, and M_reproj_ = C^r^ * M_proj_.

To verify the significance of the overlap, we permuted the order of coefficients in the columns of C_s2_ to get C_s2_. We only used C_s2_ when the total variance of C_s2_ *M was in between 0.8 - 1.2 times the total variance of C_s2_ * M. We then repeated subspace overlap calculations for 100 different C_s2_ meeting this criteria. This ensured that the resulting null distribution of overlap values did not differ from the actual value simply because the C_s2_ captured much less variance than C_s2_. We also calculated subspace overlap for subspaces that were each computed using just one half of the total epochs from the neural recording, repeating this for 300 different random parcellations of the epochs.

## Acknowledgments

The authors would like to thank J. Glaser, D. Dombeck, M. Elbaz and D. Xing for helpful advice. This research was supported by a Searle Scholar Award, a Sloan Research Fellowship, a Simons Collaboration on the Global Brain Pilot Award, a Whitehall Research Grant Award, The Chicago Biomedical Consortium with support from the Searle Funds at The Chicago Community Trust, and NIH grant DP2 NS120847 awarded to A.M.

## Author Contributions

The climbing wheel apparatus was designed and built by A.S. with guidance from A.M. Data were collected by N.K. and Z.M., with help from A.S. and A.K. Data analysis methods were developed and applied by Z.M., N.K., and A.M., with early assistance from M.A. and M.Y.

## Materials & Correspondence

Correspondence and request for materials should be addressed to A.M.

## Data and code availability

The data that support the findings of this study, all Matlab code used for data analyses, and CAD files for 3D printed wheel components will be made available on the Miri lab’s GitHub page upon publication.

---

**Extended Data Figure 1.**
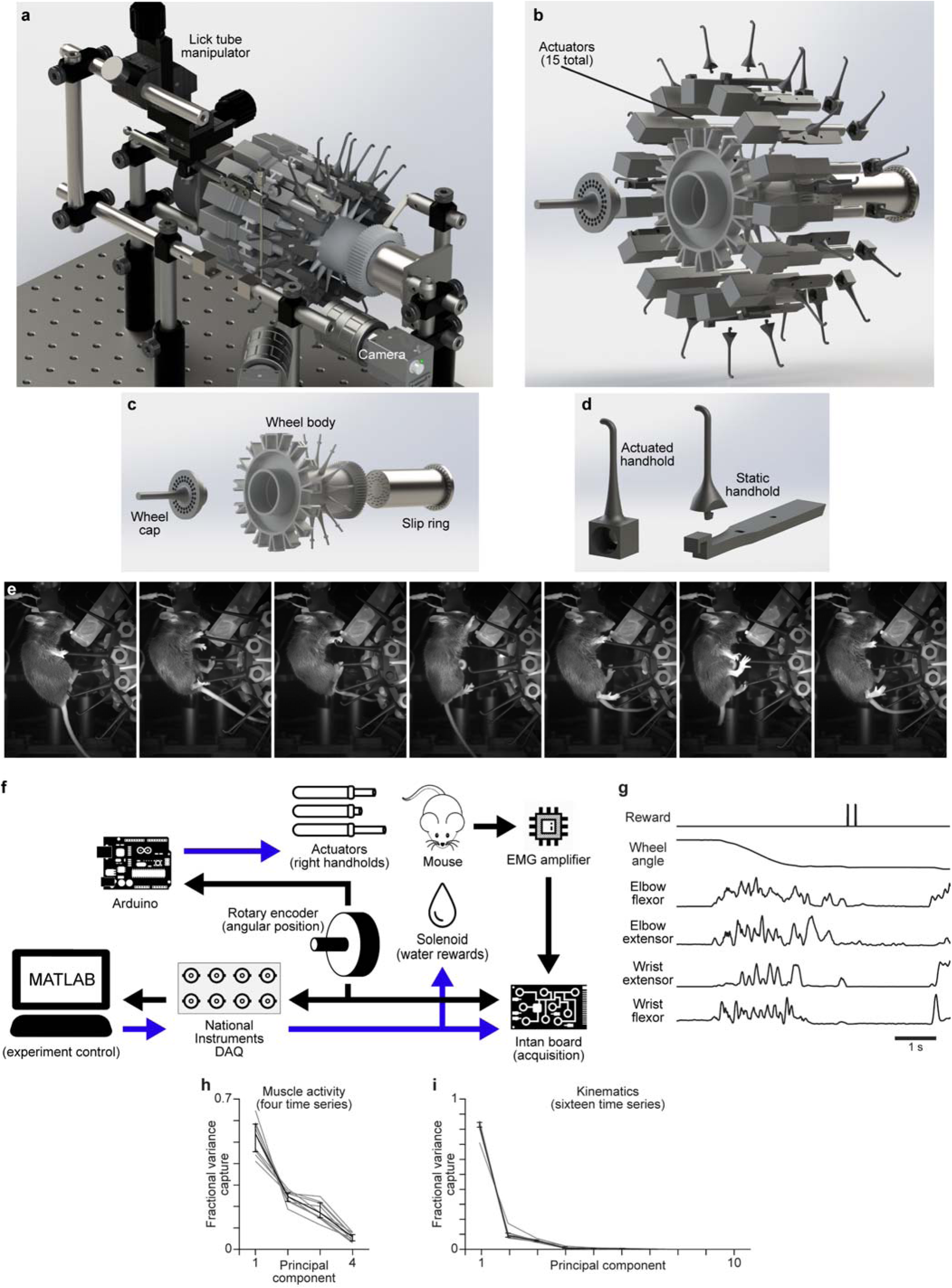
Head-fixed climbing paradigm. **a**, 3D illustration of the climbing apparatus, including the lick tube manipulator with lick tube, mounts for head fixation, and cameras. **b**, 3D exploded illustration of the wheel with the separate components visible, showing the slip ring, actuators, and 3D printed parts. **c**, Same as **b**, but without actuators and handholds. **d**, The two types of 3D printed handholds. Actuated handholds slide onto actuators. Static handholds attach to the wheel body between actuators. **e**, Seven video frames illustrating the range of different postures mice express during climbing. **f**, Flow chart illustrating data acquisition and experimental control. Blue arrows indicate command output signals. Black arrows indicate measured signals. **g**, Example time series during climbing: solenoid command for dispensing reward, wheel angle from optical encoder, and four channels of EMG. **h**,**i** Whisker plots spanning the 1^st^ to 3^rd^ quartiles (black, n = 8 mice) for the variance captured by principal components of muscle activity time series (**h**) and kinematic time series (**i**, x- and y-coordinates for eight tracked points) from one session for each mouse. Gray lines are for individual mice. Three principal components captured the vast majority of the variance for both muscle activity and limb kinematic time series. Limb orientation is charted using video-based tracking^71^.

**Extended Data Figure 2.**
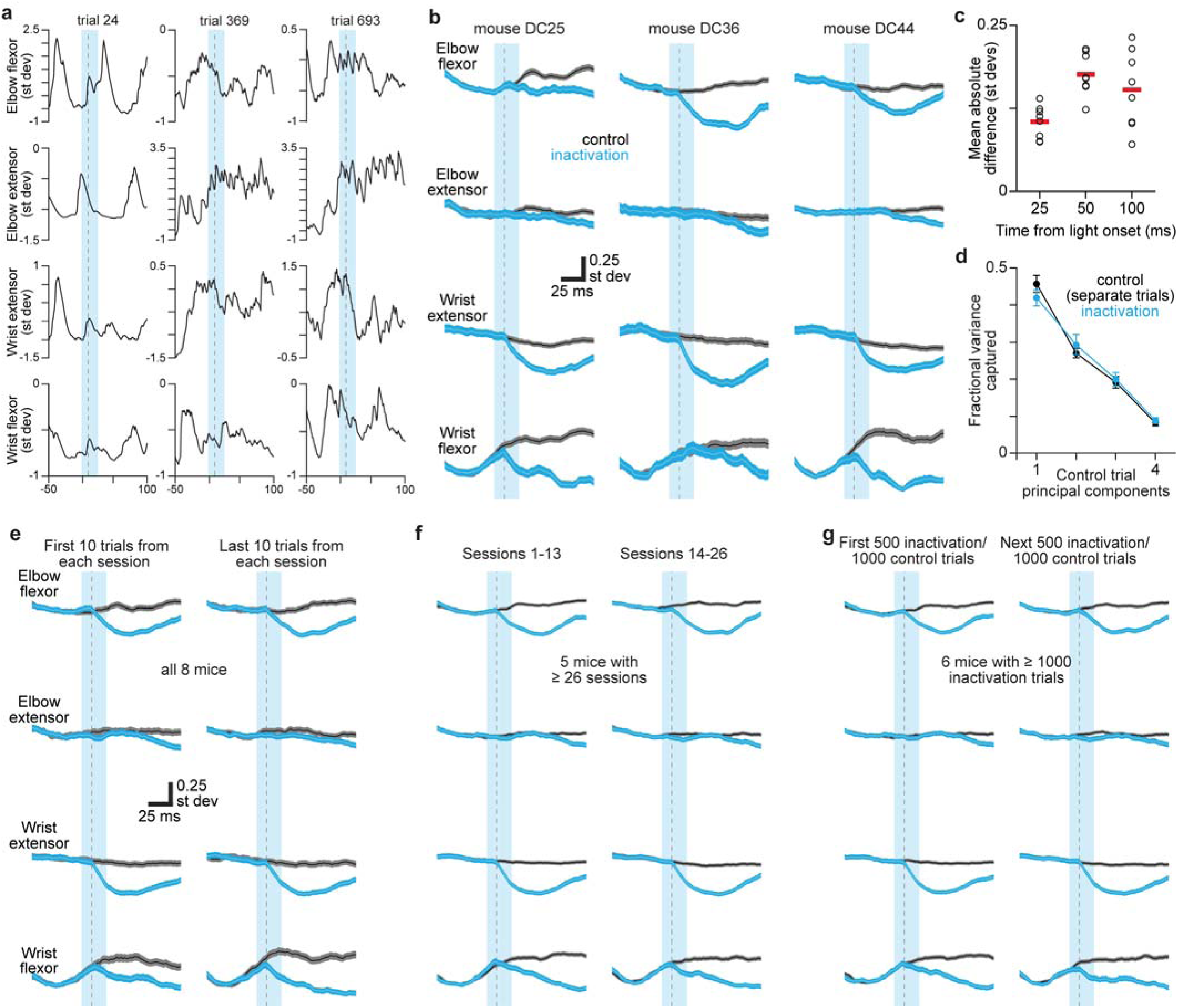
Effect of CFA inactivation on muscle activity. **a**, Muscle activity times series surrounding three inactivation trials in one mouse. The activity of each muscle was z-scored, so units are standard deviations of the original recorded signal. Vertical cyan bars in **a**,**b**,**e**-**g** indicate the 25 ms epoch of blue light applied to CFA. Gray dotted lines are 10 ms after light onset, the shortest latency at which effects can be observed. **b**, Mean ± sem muscle activity for control (gray) and inactivation (cyan) trials for the recorded forelimb muscles in three additional mice. **c**, Absolute difference between inactivation and control trials at 25, 50, and 100 ms after trial onset for individual animals (black circles) and the mean across animals (red bars). The mean was significantly greater than 0 at 25, 50 and 100 ms following light onset (p = 0.004 in each case, Wilcoxon signed-rank test). Despite this, we found that the covariance of muscle activity patterns was not substantially different in the 50 ms following control and inactivation trial onsets **d**, Mean ± sem (n = 8 mice) variance capture of muscle activity during inactivation trials and one half of control trials using principal components computed from the other half of control trials. Only time series during the 50 ms following trial onset were used here to focus on the period when inactivation effects were most apparent. The covariance of muscle activity patterns was not substantially different in the 50 ms following control and inactivation trial onsets. **e**-**g**, Mean ± sem muscle activity for control (gray) and inactivation (cyan) trials using the first 10 and last 10 trials of each type from each session (**e**), the first 13 and next 13 sessions for each mouse (**f**, n = 5 mice), and the first 500/1000 inactivation/control trials and the next 500/1000 inactivation/control trials for each mouse (**g**, n = 6 mice). Average inactivation effects on muscle activity show remarkable consistency, both within and across sessions.

**Extended Data Figure 3.**
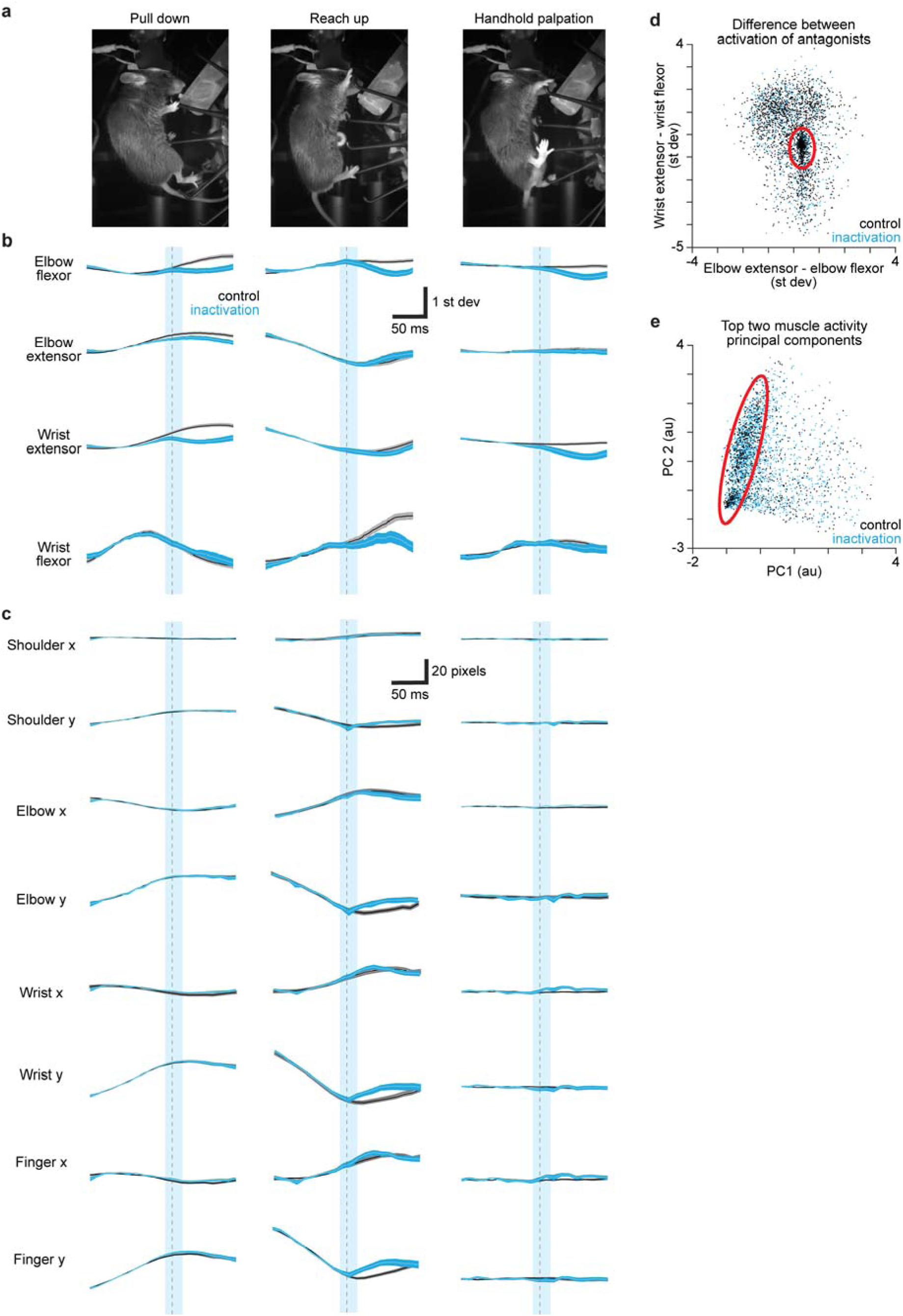
CFA inactivation effects during stereotypical features of climbing behavior. a, Extracted video frames illustrating three common behavioral features during climbing from one example animal. **b**, Trial-averaged (mean ± sem) muscle activity for control (gray) and inactivation (cyan) trials for forelimb muscles in the example animal. Vertical cyan bars indicate the 25 ms epoch of blue light applied to CFA. Gray dotted lines are 10 ms after light onset, the shortest latency at which effects can be observed. Note that in **b** and **c** the time series for palpation appear largely flat because the oscillations in muscle activity and limb position during palpation were not aligned across trials, and so much of the structure of individual trials averaged out in trial averages. **c**, Trial-averaged (mean ± sem) time series of four tracked joints for control (gray) and inactivation (cyan) trials in the example animal. **d**, For all control (black) and inactivation (cyan) trials from one representative mouse, the difference between the activation of the two wrist muscles averaged over the 50 ms prior to trial onset plotted against the corresponding difference in activation of elbow muscles. Red ellipses in **d** and **e** indicate regions where trials are highly clustered, challenging our ability to divide them into groups. **e**, For all control (black) and inactivation (cyan) trials from the same mouse used in **d**, a scatterplot of the mean projection of muscle activity over the 50 ms prior to trial onset onto the top two principal components for the activity of all four muscles.

**Extended Data Figure 4.**
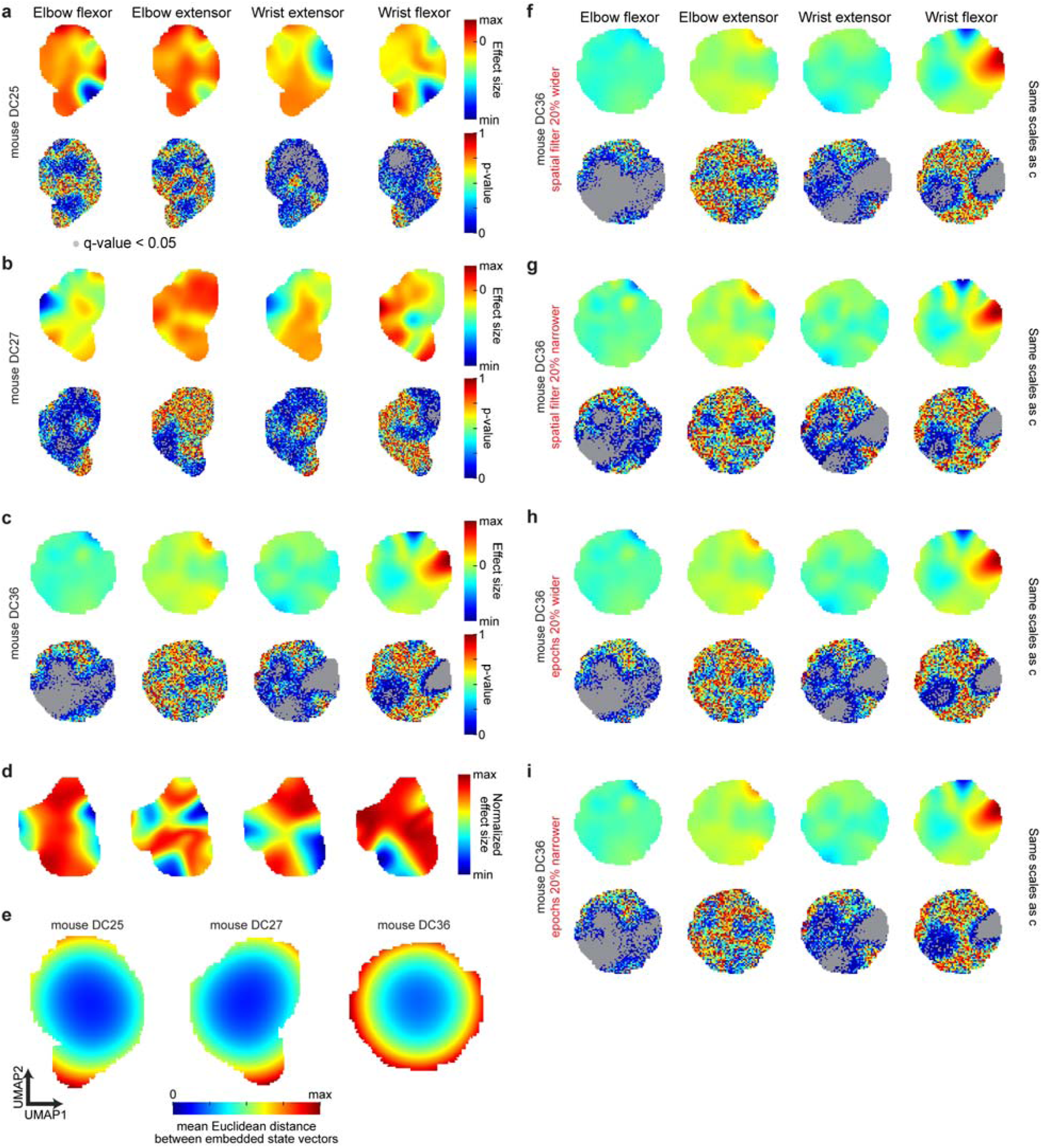
CFA inactivation effects across muscle activity state space. a-c, Inactivation effect maps and corresponding p-value maps for each of the four muscles in three additional mice. Q-values (gray overlay) reflect the expected rate of false discovery below the corresponding p-value^46^. For the effect sizes, color scale max and min reflect the maximum and minimum effect sizes across all four muscles. Thus, the maps for muscles with smaller effect sizes do not use the full range of colors. **d**, Alternate versions of the inactivation effect maps shown in Fig. 2j in which the color scale applies separately to each map, so the same colors correspond to the different maximum and minimum values on each map. **e**, For three additional mice, maps in which each grid point is colored by the mean distance (in 80D) between all pairs of embedded states, with each distance weighted by a Gaussian function of the pair’s mean distance from the grid point on the 2D map. **f-i**, Same as **c**, but following calculation with slightly different parameters (red text).

**Extended Data Figure 5.**
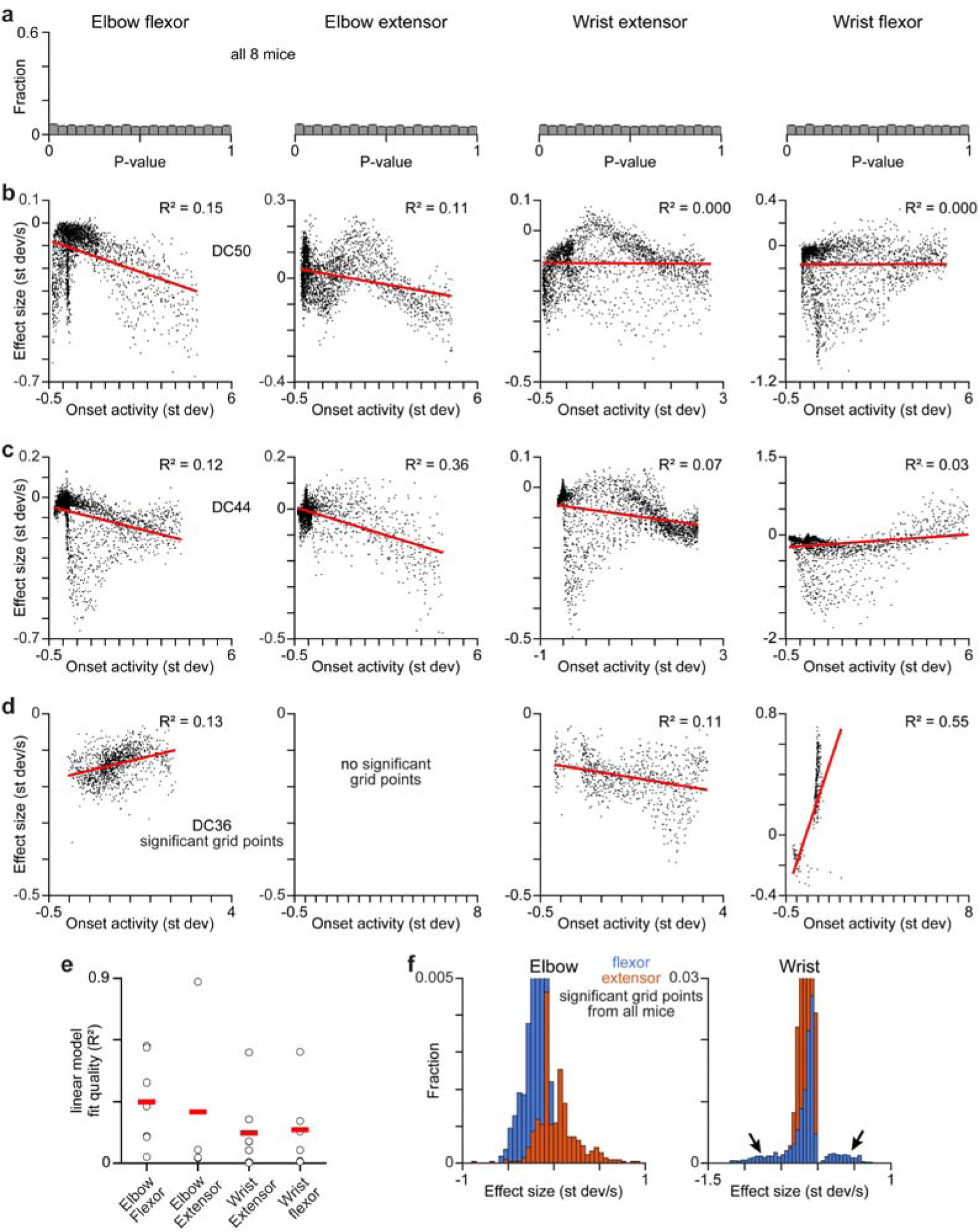
CFA selectively excites physiological flexors. **a**, P-value distributions for inactivation effects on each muscle when calculated by comparing control trials to a separate set of control trials, for all grid points across all 8 mice. **b**,**c**, Scatterplots of inactivation effect size versus muscle activity at trial onset for two additional mice. Each point reflects a different grid point. R^2^ is for a linear fit (red). **d**,**e** Same as Fig. 3f (**d**) and 3g (**e**), except only including grid points where effect size was significantly different from zero. Note that the y-axis scale in **e** differs from that in Fig. 3g. As was true when using all effect sizes, residuals from linear fits for these subsets were significantly different from uniform (p < 0.004 for all mice, K-S test). **f**, Same as Fig. 3j, but zoomed in 10-fold to clarify rare larger effects.

**Extended Data Figure 6.**
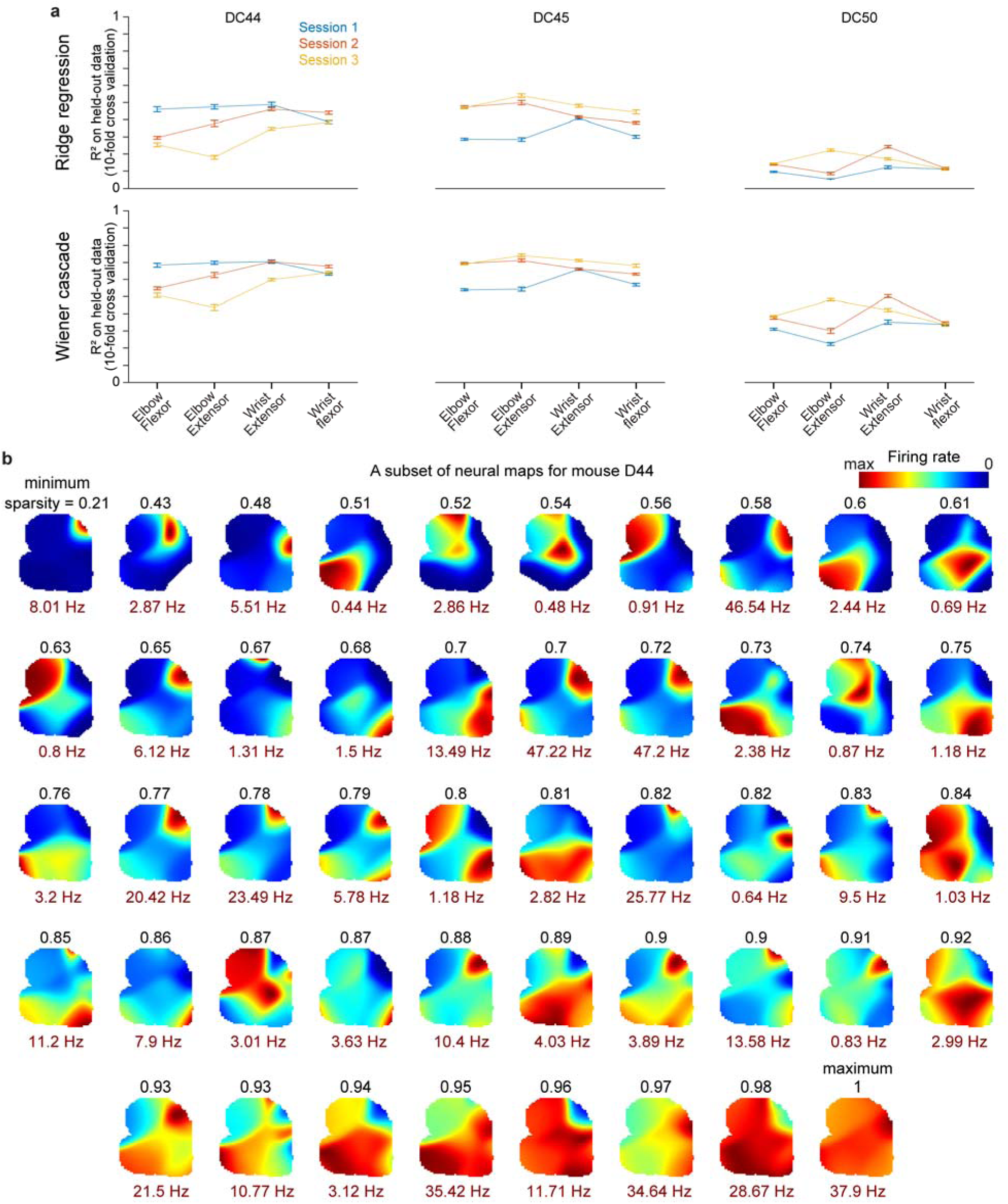
Analysis of neural recordings. **a**, For each mouse, mean ± sem (n = 10 folds) R^2^ for ridge regression (top) and Weiner cascade^73^ (cubic nonlinearity, bottom) models that fit muscle activity with the top principal components (PCs) capturing at least 99% of variance in neuronal firing rates. Muscle activity averaged in 10 ms bins was fit using the neural PCs over the preceding 100 ms (10 bins). The middle 50% of each recording was used to reduce compute time. This segment was divided into 1-second epochs, and a random 10% of epochs were held out for performance measurement. The first 100 ms of these epochs were omitted from testing to avoid overlap with training data. The mean ± sem R^2^ values across sessions and muscles were 0.310 ± 0.051 (ridge) and 0.546 ± 0.051 (Wiener). **b**, Neural activity maps for 48 neurons from one mouse. To illustrate varying sparsity, neurons shown have values equally spaced along the full distribution of sparsity values for the given mouse. Black text is sparsity, red text is each map’s maximum firing rate.

**Extended Data Figure 7.**
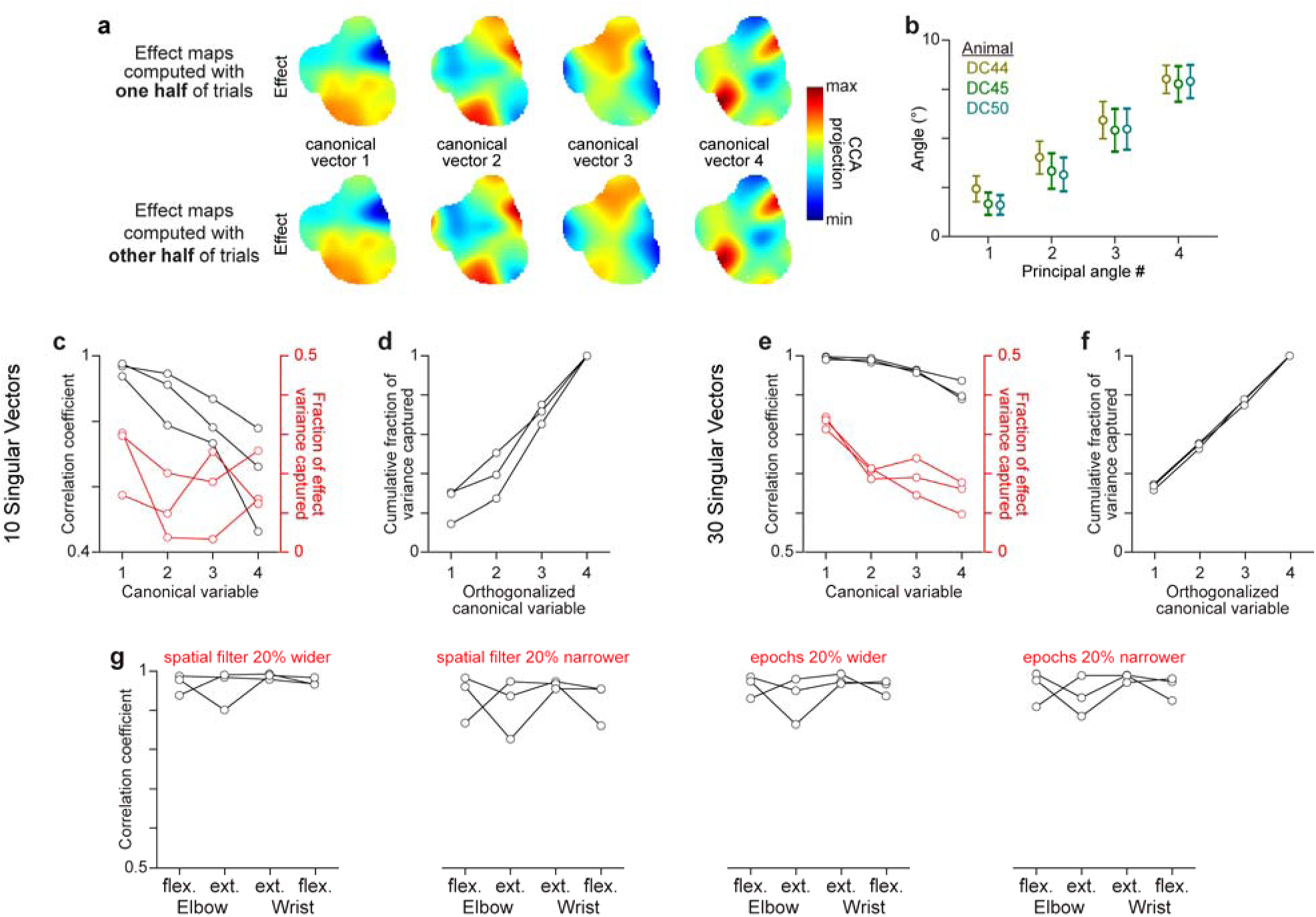
Finding a CFA activity subspace that aligns with CFA influence. **a**, Canonical variables for inactivation effect maps resulting from CCA with neural activity maps, using effect maps computed with separate halves of trials. **b**, Mean ± standard deviation for principal angles between subspaces spanned by the neural canonical vectors from CCA that used effect maps computed with separate, randomly-chosen halves of trials, over 300 iterations. **c**,**d** Correlation coefficient (black) and fractional inactivation effect variance captured (red) for canonical variables (**c**), and cumulative fraction of effect variance captured by canonical variables after orthogonalizing their corresponding vectors (**d**) when SVD is used to reduce neural activity dimensionality to 10 instead of 20 before applying CCA. Each set of connected dots in **c-g** is from one animal. **e**,**f**, Same as **c**,**d**, but when SVD is used to reduce neural activity dimensionality to 30. **g**, Correlation coefficients from CCA aligning neural activity maps and inactivation effect maps for individual muscles, calculated after changing key inactivation map parameters (red text).

**Extended Data Figure 8.**
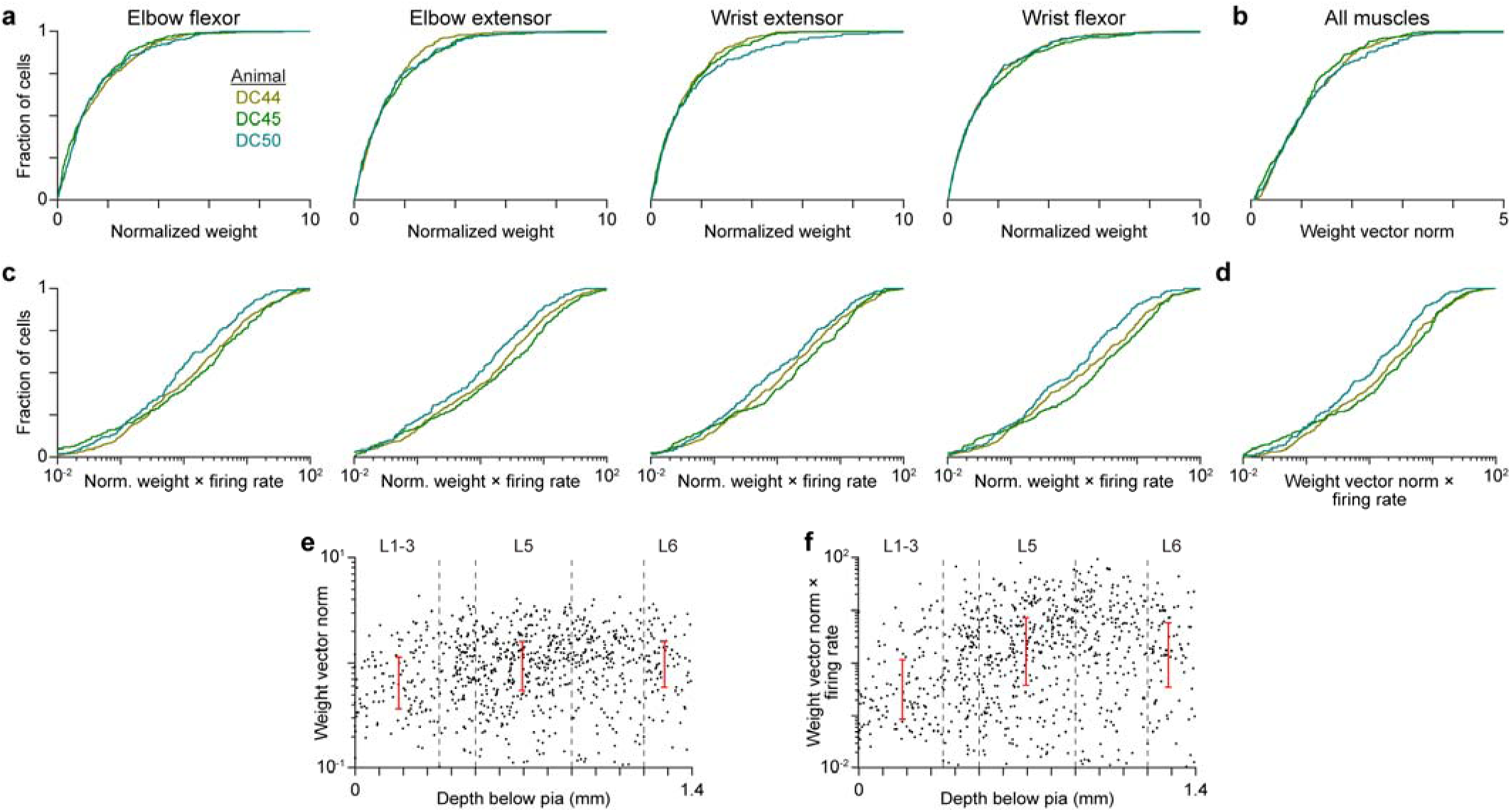
Influence subspace contributions across neurons. Here used the canonical vectors defined using the inactivation effect maps of individual muscles. **a**, Cumulative distributions of weights for the activity of individual neurons in canonical vectors defined by CCA using the inactivation effect maps of individual muscles. Weights were normalized so the median value for each mouse is 1 to enable comparison across mice, as different numbers of neurons were recorded in each mouse. Tests for multimodality were not significant for all distributions (p-values ranged from 0.64 to 0.99, Hartigan’s dip test). Results indicate a broad distribution of neuronal weights for each muscle in each mouse. **b**, Cumulative distribution of norms of the four-element vectors composed of each given neuron’s canonical vector weight from alignment with the inactivation effect maps of individual muscles. Tests for multimodality were not significant for all distributions (p-values ranged from 0.34 to 0.99, Hartigan’s dip test). Results again indicate a broad distribution of neuronal weights for each muscle in each mouse. **c**,**d** Same as **a**,**b**, but weights (**c**) and weight vector norms (**d**) are scaled by each neuron’s mean firing rate during climbing. Tests for multimodality were not significant for all distributions (p- values ranged from 0.90 to 0.99, Hartigan’s dip test). **e**, Scatterplot of weight vector norms as in **b** versus the depth below pia assigned to the waveform centroid of each neuron. Red whisker plots span from the first to the third quartile of neurons assigned to each laminar group (those within dotted boundaries). Data from three mice are combined. The distributions of weight vector norms for neurons localized to layers 5 (p = 3 x 10^-5^, Wilcoxon’s rank sum test) and 6 (p = 7 x 10^-4^) were significantly higher than those localized to superficial layers. **f**, Same as **e**, but weight vector norms are scaled by each neuron’s mean firing rate during climbing. The distributions of weight vector norms for neurons localized to layers 5 (p = 5 x 10^-14^, Wilcoxon’s rank sum test) and 6 (p = 3 x 10^-7^) were significantly higher than those localized to superficial layers.

**Extended Data Figure 9.**
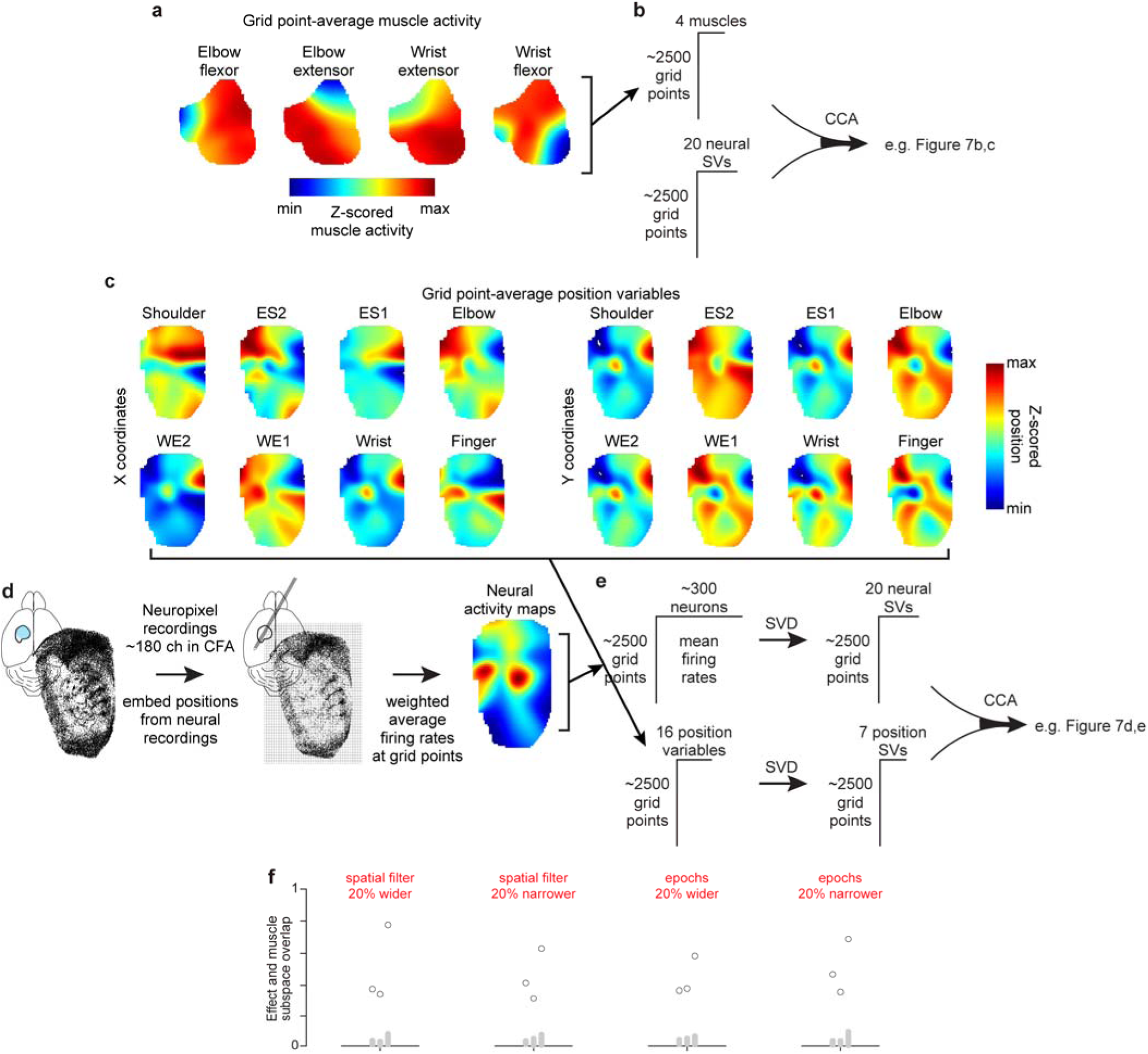
Extracting neural activity subspaces aligned with muscle activity and limb kinematics. **a**, Grid point-averaged muscle activity maps for an example animal. **b**, Grid point-averaged muscle activity maps are used to create a grid points by muscles matrix that, together with a grid points by 20 neural singular vectors matrix, serve as inputs to CCA. CCA then yields the canonical variables illustrated in Figure 7b,c. **c**, Grid point-averaged position maps for the eight tracked sites along the right forelimb for an example animal. **d**, 50 ms segments of the site positions tracked during neural recording sessions are embedded into the 2D limb orientation state map, and the neural activity corresponding to the embedded vectors is used to calculate grid point-averaged neural activity maps. **e**, The maps in **c** can be collapsed into a grid points by position variables matrix which is then dimensionally reduced to a grid points by 7 position singular vectors matrix using SVD. This matrix and the grid points by 20 neural singular vectors matrix obtained from the neural activity maps serve as input to CCA. CCA then identifies the canonical variables shown in Figure 7d,e. **f**, Overlap between the influence and muscle activity subspaces (black circles) compared to 300 estimates of the overlap expected by chance (gray dots) for each animal calculated after slight changes to the parameters used for map calculations (red text).

**Extended Data Figure 10.**
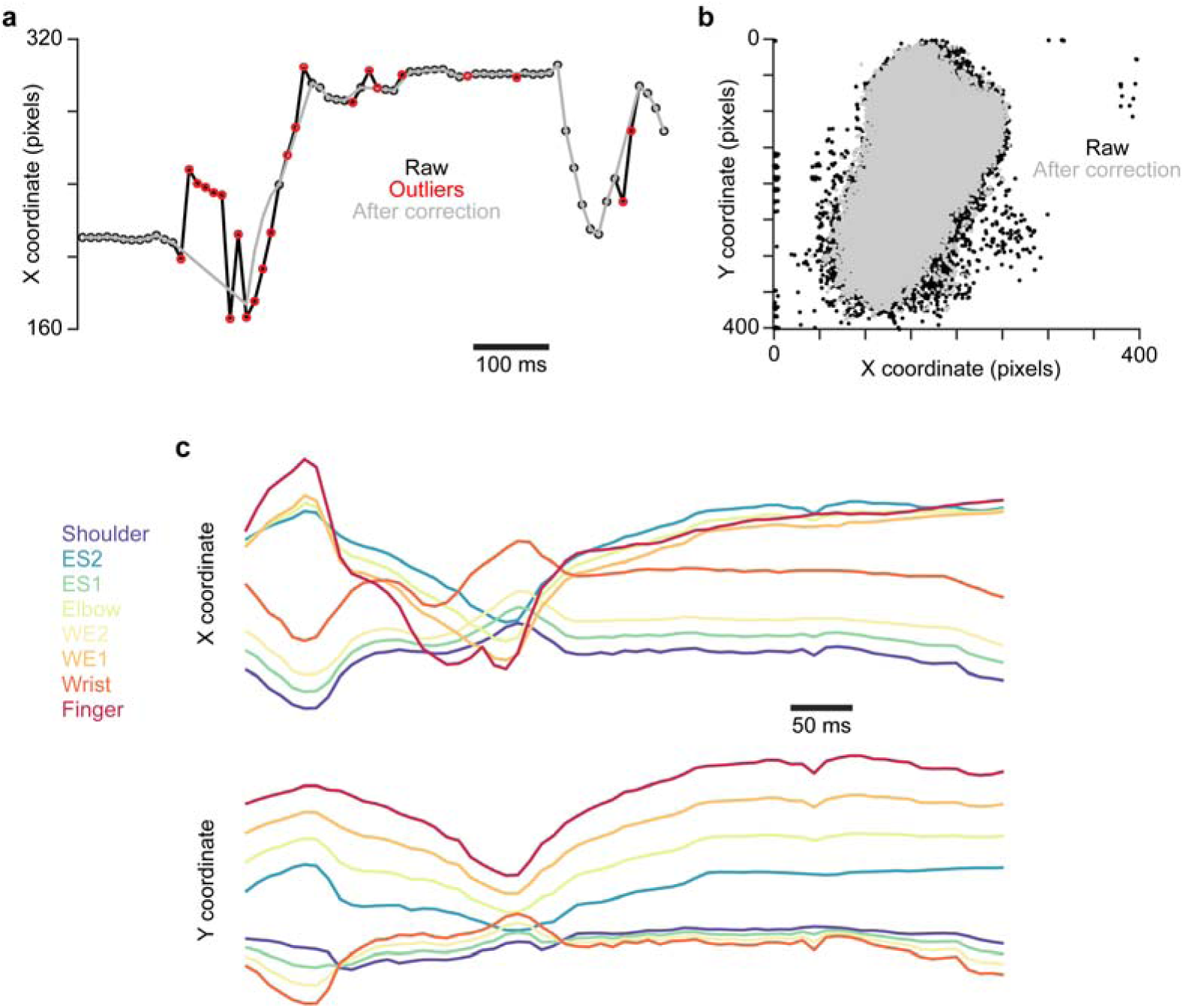
Post-processing of video-based limb tracking. **a**, Example time series for the point on the finger, illustrating post-hoc processing to correct outliers. Raw is the time series output by DeepLabCut before correction. **b**, Scatter plot of X and Y coordinates of the finger site tracked in **a**, showing fewer outliers after correction. **c**, Example X and Y coordinate traces for site positions (defined in Fig. 4) after post-hoc processing.

**Supplementary Movie.**
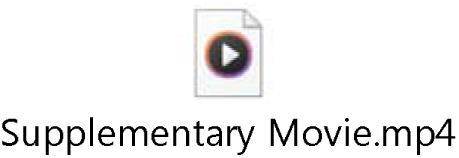
Head-fixed climbing. An example of a mouse climbing in our paradigm. The bird’s-eye view towards the end makes apparent the stagger of the handholds on the mouse’s right; this stagger itself is ever changing and thus unpredictable.

